# Cholesterol metabolism is a potential therapeutic target in Duchenne Muscular Dystrophy

**DOI:** 10.1101/2020.12.01.405910

**Authors:** F. Amor, A. Vu Hong, G. Corre, M. Sanson, L. Suel, S. Blaie, L. Servais, T. Voit, I. Richard, D. Israeli

## Abstract

**Background:** Duchenne Muscular Dystrophy (DMD) is a lethal muscle disease detected in approximately 1:5000 male births. DMD is caused by mutations in the DMD gene, encoding a critical protein that link the cytoskeleton and the extracellular matrix in skeletal and cardiac muscles. The primary consequence of the disrupted link between the extracellular matrix and the myofiber actin cytoskeleton is thought to involve sarcolemma destabilization, perturbation of Ca^+2^ homeostasis, activation of proteases, mitochondrial damage and tissue degeneration. A recently emphasized secondary aspect of the dystrophic process is a progressive metabolic change of the dystrophic tissue; however, the mechanism and nature of the metabolic dysregulation is yet poorly understood. In this study, we characterized a molecular mechanism of metabolic perturbation in DMD.

**Methods:** We sequenced plasma miRNA in a DMD cohort, comprising of 54 DMD patients treated or not by glucocorticoid, compared to 27 healthy controls, in three age groups. We developed an original approach for the biological interpretation of miRNA dysregulation, and produced a novel hypothesis concerning metabolic perturbation in DMD. We then used the mdx mouse model for DMD for the investigation of this hypothesis.

**Results:** We identified 96 dysregulated miRNAs, of which 74 were up- and 22 down-regulated in DMD. We confirmed the dysregulation in DMD of Dystro-miRs, Cardio-miRs and a large number of the DLK1-DIO3 miRNAs. We also identified numerous dysregulated miRNAs, yet unreported in DMD. Bioinformatics analysis of both target and host genes for dysregulated miRNAs predicted that lipid metabolism might be a critical metabolic perturbation in DMD. Investigation of skeletal muscles of the mdx mouse uncovered dysregulation of transcription factors of cholesterol and fatty acid metabolism (SREBP1 and SREBP2), perturbation of the mevalonate pathway, and accumulation of cholesterol. Elevated cholesterol level was also found in muscle biopsies of DMD patients. Treatment of mdx mice with Simvastatin, a cholesterol-reducing agent, normalized these perturbations and partially restored the dystrophic parameters.

**Conclusion:** This investigation supports that cholesterol metabolism and the mevalonate pathway are potential therapeutic targets in DMD.

## Introduction

Duchenne muscular dystrophy (DMD) is the most common inherited pediatric muscle disorder. It is an X-linked genetic progressive myopathy characterized by muscle wasting and weakness, which leads to loss of motor functions, cardiac and respiratory involvement, and premature death ^1, 2^. DMD occurs at a rate of approximately 1:5000 male births and arises due to mutations in the dystrophin gene. The disease is caused by a deficiency of functional dystrophin, a critical protein component of the dystrophin glycoprotein complex acting as a link between the cytoskeleton and the extracellular matrix in skeletal and cardiac muscles ^3^.

The only routinely used medication for DMD patients is glucosteroid drugs, which can at best only slightly delay the progression of the disease ^4^. Experimental therapeutic approaches, based on gene therapy, cell therapy and drug discovery, are focused on the restoration of dystrophin expression (review in ^5, 6^). However, despite increasing efficiency in the restoration of dystrophin expression in recent clinical trials, only modest muscle functional improvement has been achieved ^7–9^.

The primary direct consequence of the disrupted link between the extracellular matrix and the myofiber actin cytoskeleton due to lack of dystrophin is thought to involve sarcolemma destabilization, perturbation of Ca^+2^ homeostasis, activation of proteases, mitochondrial damage and tissue degeneration. The tissue damage activates a regenerative response, resulting in repeated cycles of myofiber degeneration and regeneration. This ongoing process evolves into a cascade of downstream pathological events, including chronic inflammation, oxidative damage, and the replacement of contractile tissue by fibrotic and fatty tissues ^10^. A gradual decline in the capacity of the stem cells for a compensatory proliferation, differentiation, and tissue regeneration ^11^ participates also in the pathophysiology of the disease. A recently emphasized aspect of the dystrophic process is a progressive metabolic change of affected muscle^12, 13, 14–21, 22, 23^.

Profiling miRNA can be useful for diagnosis, monitoring and understanding mechanisms of diseases. DMD has been the subject of a large number of miRNA profiling studies ^24, 25, 26–33, 34–36^. However, these studies have been hampered by one or a combination of limitations, including the use of animal models rather than human patients, the relatively small size of the studied cohorts, the detection of a small number of predefined miRNAs, or the use of miRNA profiling technologies with low sensitivities. In the present study, we overcame these limitations by profiling circulating miRNAs in the plasma of a relatively large patient cohort, which was composed of three age subgroups between the ages of 4 and 20 years old, and included both glucocorticoids treated and untreated DMD patients, as well as an age-match control group. Additionally, the profiling technology was of miRNA sequencing, maximizing the detection sensitivity of the entire spectrum of expressed miRNAs. Finally, an original bioinformatics model for the interpretation of miRNA dysregulation was employed.

Interestingly, the bioinformatics analysis of plasma miRNAs dysregulation identified lipid metabolism as the most important metabolic perturbation in DMD. In order to validate this prediction, we screened muscle biopsies of young mdx mice for the expression of factors that are related to lipid metabolism, and identified their dysregulation as early as at the age of 5- weeks. Specifically, the mevalonate pathway that controls the synthesis of cholesterol was highly affected and cholesterol level was increased in the dystrophic muscles. A recent paper reported a positive effect of simvastatin on the dystrophic parameters and muscle function in the mdx mouse. Surprisingly, considering the known target of simvastatin, its effect was reported to be unrelated to blood cholesterol ^37^. We treated mdx mice with simvastatin and confirmed improved dystrophic muscle parameters. We demonstrated that components of the muscle mevalonate pathway and cholesterol levels were dysregulated in the muscle of mdx mouse compared to those of the healthy control, and returned to normal after simvastatin treatment. In addition, the normalization of lipid metabolism correlates with improved dystrophic parameters in mdx dystrophic muscle. In conclusion, besides to the discovery of a large number of dysregulated miRNA in the plasma in DMD patients, the present study provides new understanding of the metabolic perturbation in DMD and thus, opens up new perspectives for the treatment of DMD.

## MATERIALS AND METHODS

### Ethical declaration

The human study (DMD patients and controls) was conducted according to the principles of the declaration of Helsinki “ethical principles for medical research”, and was specifically approved by the ethical committee CPP Ile de France VI, on July 20, 2010, and the Comité d’Ethique (412) du CHR La Citadelle (Liège, Belgium) January 26, 2011.

### Human patients and cohort composition

DMD patients were admitted from 10 European medical centers, two of them from Belgium (n=35), one from Romania (n=16), and seven from France (n=49). Healthy control patients were admitted from the same medical centers, Belgium (n=28), Romania (n=45) and France (n=50). Patients were divided into three age groups 4-8, 8-12, and 8-20 years old. In the youngest age group, the same donors contributed the GC-treated and untreated samples, with untreated samples that were obtained before, and treated samples after their first GC treatment, at interval of less than 6 month. GC-treated and untreated samples for the age groups of 8-12 and 12-20 years old were obtained from distinct DMD patients.

Human skeletal muscle tissues were obtained from the Myobank, the tissue bank of the Association Francaise contre les Myopathies (AFM). Open skeletal muscle biopsies were performed after informed consent, according to the Declaration of Helsinki. Muscle biopsies included in this study were derived from the paravertebral (two controls and two DMD), Gluteus, and the tensor fasciae latae (one each control and one DMD) from the group aged 8-12 (DMD) and 12-20 (control) years old.

### Blood collection

Samples were collected from individuals with a written informed consent of parents or legal guardians. Human blood samples were collected from male subjects of at least 3 years-old and 15kg body mass, in both control and DMD patients. Peripheral blood samples were collected into 5 ml K3EDTA tubes (Greiner Bio-One). Plasma was separated from buffy coat and red blood cells after 10 minutes centrifugation at 1800 g and stored at −80°C until further processing.

### Choice of RNA Samples for the miRNA profiling

Following the RNA extraction, a sub-cohort of 81 RNA samples was selected for the high throughput sequencing (HTS) cohort (**Supplemental table 2**). We employed an optimization process for the selection of the most suited RNA samples. This included the selection of (i) plasma samples of OD 414 > 0.2 (indicative of absence of hemolytic contamination ^38^, (ii) removal of RNA preparation of low concentration (< 30 ng/µl, conferring improved RNA stability), (iii) samples with optimized patients’ age-distribution (inside age-groups), and (iv) age-matching between group types (i. e. between DMD, DMD non treated and healthy controls). Thus, the entire studied HTS cohort was composed of 54 DMD patients and 27 healthy controls (N=81) divided into 9 groups of 9 subject, composed of three age groups as described in **Figure 1A.**

**Figure 1.**
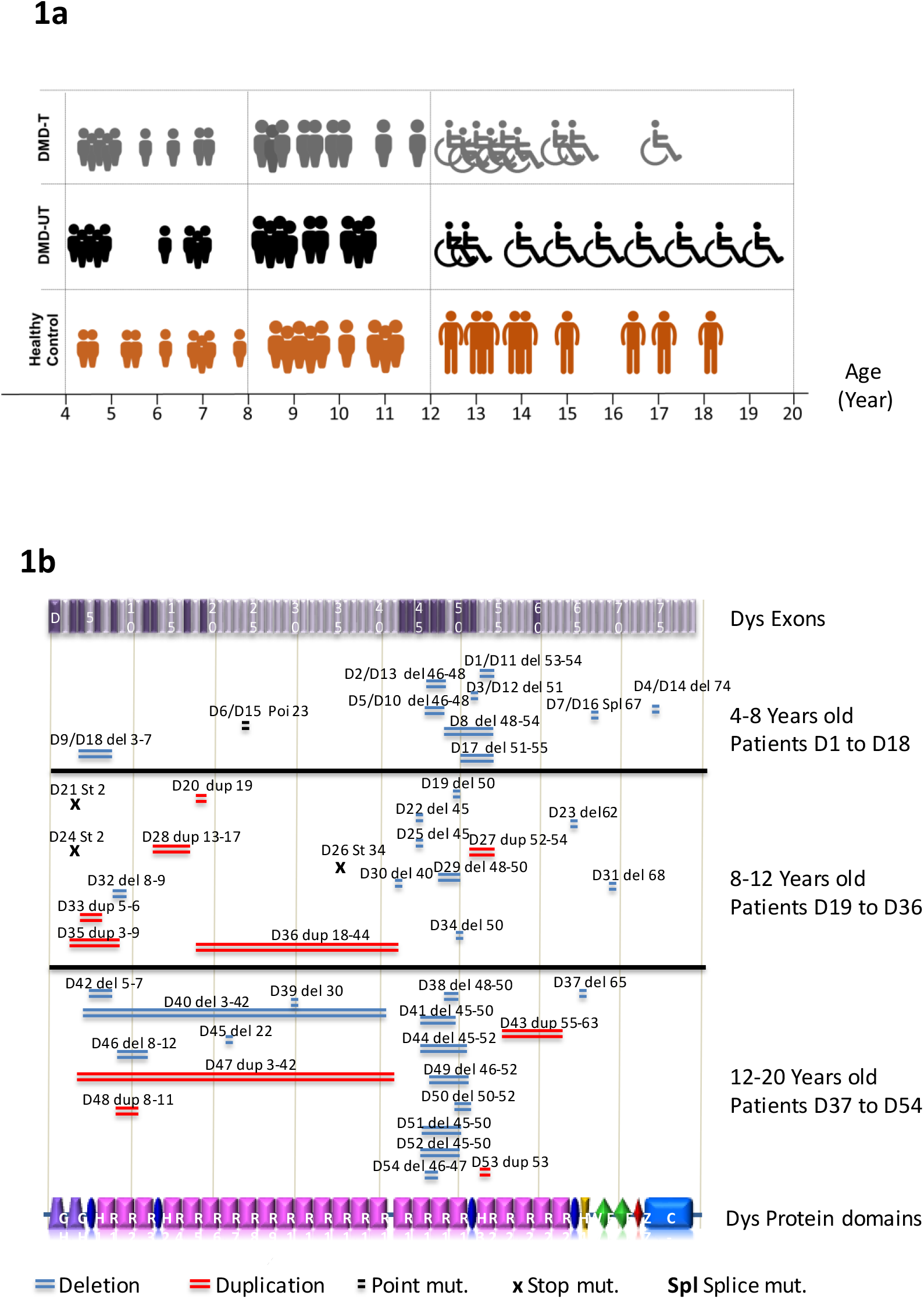
DMD cohort characterization. (**1a)** Cohort subgroups and age composition. DMD patients and healthy controls were classified into three age groups of 4-8, 8-12 and, 12-20 years old. DMD patients were glucocorticoid treated (DMD T) or untreated (DMD UT). The figurine symbols represent the ages of individual patients on the horizontal axis, black for untreated DMD, grey figures for treated DMD, and brown figures for healthy controls. Treated and untreated DMD of the 4-8 group of age are the same patients before and after GC treatment (except one patient). **(1b)** A graphical presentation of the spectrum of dystrophin mutations by age group. Dystrophin’s gene 79 exons and protein domains are presented on respectively the upper and lower vertical bands. Patients of the 4-8 age years old group are represented twice in the cohort (with the exceptions of D8 and D17), before and after Glucocorticoid treatment, by samples D1 to D9 and D10 to D18 respectively. Del (blue) =deletion; Dup (red) = duplication; St (black) = stop codon mutation; Poi (in black) = point mutation.

### MiRNA Sequencing

MiRNA sequencing was performed by Integragen (Evry, France). Libraries cloning was modified from ^39^, for improved efficiency in small samples. Briefly, a 3’ adenylated DNA adaptor was ligated in the presence of 12% PEG and the absence of ATP, avoiding miRNAs self-ligation. A 5’ RNA adaptor was ligated in the presence of ATP. RT primer complementary to the 3’ adaptor was added, forming a duplex to reduce adapter dimer formation. RT reaction was carried out with 1.75 pmol adaptors (3’ adaptor / 5’ adaptor / RT primer), and cDNAs were amplified by 13 PCR cycles with primers complementary to the 3’ and 5’ adaptors. During this PCR step, a specific barcode was incorporated for individual sample recognition. PCR samples band quantification was carried out with Fragment Analyzer (AATI). An equimolar pool of ten different samples was migrated on PAGE, and the miRNA band was extracted (Qiagen MinElute column). Libraries were quantified by qPCR, to load precisely 7pM pool per line of HiSeq Flow-Cell. The HiSeq 36b and index (barcode) sequencing was carried out as instructed (Illumina) with a SBS V3 kit leading on 150 million passing filter clones.

All clean reads were compared to the Rfam database (http://rfam.xfam.org/), repeatmasker (http://www.repeatmasker.org/, UCSC download 01/04/2014), and the NCBI RefSeq ^40^, download 10/04/2014) for the annotation of the rRNA, snoRNA, piRNA ^41^, and tRNA ^42^. Unique miR reads and their copy numbers were aligned with miRanalyzer online software ^43^, using Ensembl human gene browser (genome assembly GRCh38) and (miRbase v20, June 2013 ^44^). MiR count raw data was normalized and processed for differential expression by the Deseq2 R package ggplot2 ^45^. The significance of miRNAs differential expression was ranked by T-test, with a false discovery rate (FDR) correction according ^46^.

### MicroRNA target genes enrichment analysis

MiR-target gene enrichment analysis was performed with the online mirPath v.3^47^ http://snf-515788.vm.okeanos.grnet.gr/, based on predicted miRNA to mRNA interaction in human.

### MicroRNA host genes analysis

For this analysis we considered all the dysregulated miRNAs p≤0.05 in DMD patients of the aged 4-12 years old, compared to their age related healthy controls **(Supplemental table 2)**. MicroRNA host genes were retrieved from the miRIAD database ^48^ and were manually validated on the Ensembl human genome browser GRCh38.p7. We considered all miRNAs embedded on the sense strand of introns and exons of protein coding genes. We assigned to the host-genes the fold change (FC) and p values of their embedded miRNAs. When both 3p and 5p isoform of the same premiRNA were dysregulated their host-genes were considered only once, with the lower p value miR isoform. Similarly host genes for a number of different miRNAs in a miRNA cluster was considered only once, with the highest dysregulated miRNA, (according p value). In miRNAs hosted on more than one host-gene, both host-genes were considered. The table of host-genes with their assigned dysregulation values is in Supplemental table 8. Host genes data were assigned to the core analysis of Ingenuity® Pathway Analysis (IPA®, QIAGEN Redwood City,www.qiagen.com/ingenuity). Pathway enrichment of the gene set was performed by ReactomePA and ClusterProfiler R/Bioconductor packages ^49^.

## RESULTS

### DMD patient characterization and cohort composition

To perform miRNA profiling in the serum of DMD patients, we collected samples of 100 DMD patients and 123 healthy controls from 10 European medical centers **(Supplemental table 1**). The study was approved by central and local Institutional Review Board (IRB), and recorded on clinicaltrials.gov (NCT NCT01380964). All patients or parents for minors signed an informed consent. Following RNA extraction from the plasma and quality control validation, we selected 81 RNA samples and constituted nine groups of nine patients. The groups consist of DMD patients, either untreated (DMD-UT) or receiving treatment with glucocorticoid (DMD-T) and healthy controls, in three age categories of 4-8, 8-12 or 12-20 years old. (**Figure 1a).** The mean age of disease onset of the corresponding cohort was 3 years of age. Normal walking capacity was preserved in all patients of the first age group while three patients of the second group (33%) and all but 2 in the last age group (88%) had lost their ability to walk **(Supplemental table 2)**. Overall, the mean age of loss of walking ability was 9.6 years. Additional clinical and functional characterization of the patients is shown in **Supplemental table 2** and the spectrum of dystrophin mutations present in the patients is depicted in **Figure 1b**.

**Table 1:** Network analysis of host genes for dysregulated miRNAs. A pathway analysis was used for the construction of gene networks, using the core analysis of Ingenuity® Pathway Analysis (IPA) with a threshold of 70 molecule / network. The constructed networks are composed of host-genes for dysregulated miRNAs (p<0.05) in the plasma of the 4-12 years old DMD patients. The P value denotes the “probability” of the coincidental construction of a network out of its components. The Focus molecule (3^rd^ column) denotes the number of the network’s principal molecules, and the identity of the most connected molecule of each network is provided in the 4th column. The 4rth network was obtained by the merging all networks function of IPA algorithm.

|  | Network | Score | Number Focus Molecules | Most connected molecule |
| --- | --- | --- | --- | --- |
| 1 | Lipid Metabolism,<br>Small Molecule Biochemistry<br>Molecular Transport | 1 e-64 | 31 | SREBF1 |
| 2 | Lipid Metabolism,<br>Small Molecule Biochemistry,<br>Molecular Transport | 1 e-41 | 22 | Beta estradiol<br>(Estrogen) |
| 3 | Lipid Metabolism,<br>Small Molecule Biochemistry,<br>Dermatological Diseases<br>and Conditions | 1 e-32 | 18 | HNFA4 |
| 4<br>(Merged 1-3) | Lipid Metabolism<br>Molecular transport |  |  | SREBF1 |

### RNA profile dysregulation in the plasma in DMD patients is age-dependent

After size selection, the 81 RNA samples were all individually sequenced using the Illumina technique. All mapped reads were matched to the hg19 (GRCh38) human genome assembly and were assigned to RNA classes, including rRNA snoRNA, piRNA tRNA, and microRNA (miRbase v20, June 2013 ^44^) (**Figure 2a)**. We obtained 9.37, 3.68, and 2.81 million sequences on average per studied sample for total high quality (high quality-Reads), human genome mapped (Mapped reads), and mature miRNA sequences, respectively (**Figure 2b**). A Principal component analysis (PCA) was used for the identification of overall miRNA profile dysregulation between the different cohort subgroups. PCA failed to separate DMD from control samples (**Figure 2C upper left).** In contrast, PCA primary and secondary components separated the DMD (orange dots) from the control samples (blue dots) of the 4-8 and 8-12 age groups **(Figure 2C, upper middle and right respectively)**, but not of the 12-20 age group **(bottom left)**. In the 4-12 years old group (combined 4-8 and 8-12 groups), DMD patients were separated from healthy controls (**bottom middle).** However, PCA failed to separate GC-treated from untreated DMD patients **(bottom right**). These results indicate a robust disease effect, but not such a robust GC effect, on circulating miRNA in DMD patients below the age of 12. This effect is age-dependent as it is no longer observed beyond the age of 12.

**Figure 2.**
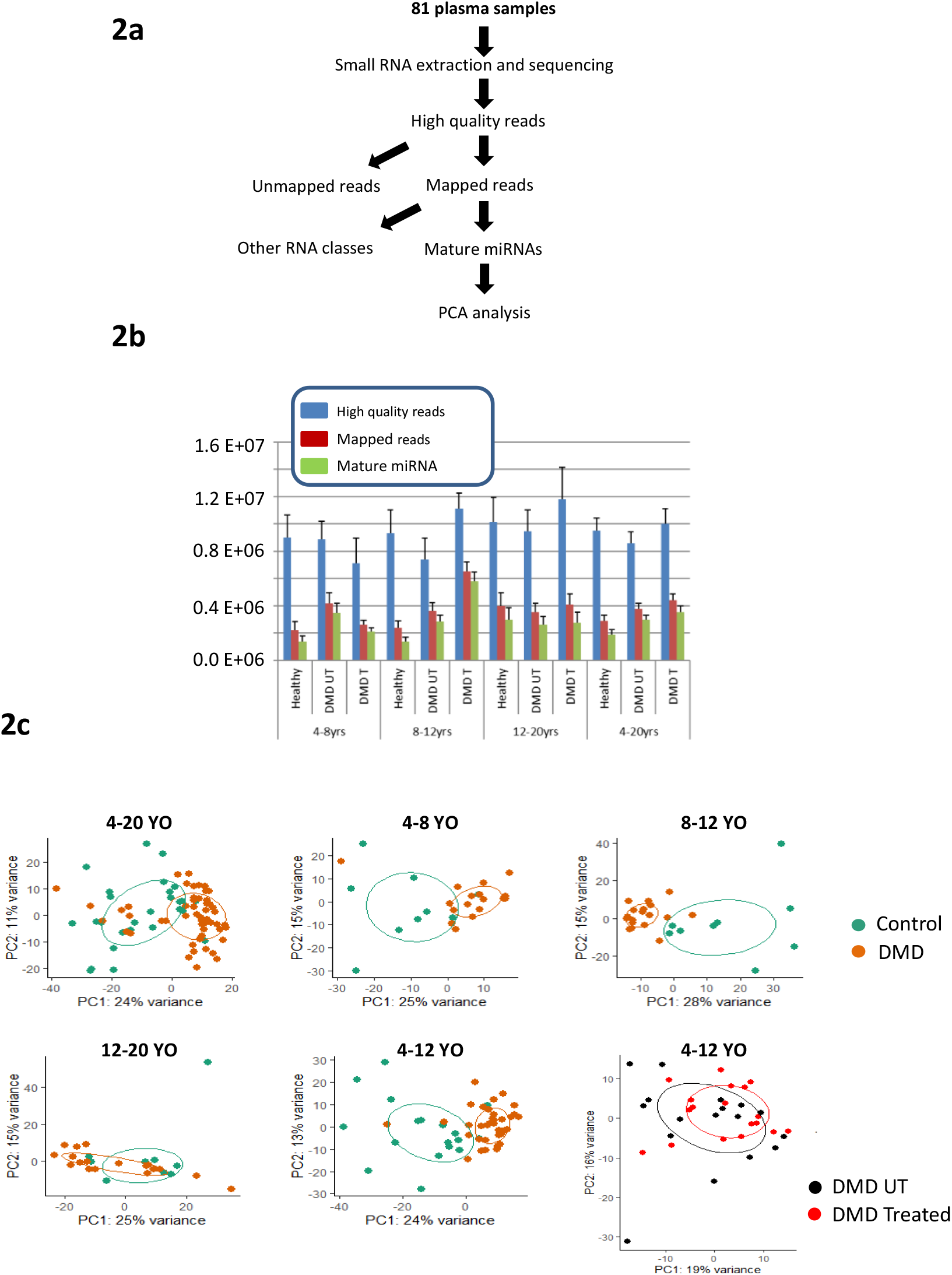
Characterization circulating RNA by high throughput sequencing. **(2a)** Schematic presentation plasma sample processing. Small RNA high quality reads (high quality Reads) were mapped on the human genome (Mapped reads). Mapped reads were classified to miRNA and other small RNA classes. **(2b)** Graphical presentation high quality Reads (blue), Mapped reads (red), and miRNA (green) in the cohort subgroups are presented as average ± SEM. **(2c)** PCA analysis of cohort segregation according to miRNA expression in DMD (orange dots) and healthy control (green dots) by age groups (panels 1 to 5) and treated (red dots) versus untreated (black dots) DMD patients (panel 6).

### Differentially expressed plasmatic miRNAs in DMD patients

The above data indicates that the overall miRNA dysregulation in DMD patients is drastically reduced beyond the age of 12. In accordance, upon stratification on age and treatment, we noticed that the largest set of dysregulated miRNAs was identified by comparing the combined GC-treated and untreated 4-12 years old patient group to their age related healthy controls (**Supplemental figure 1**), including 65 up and 25 down-regulated miRNAs (FDR<0.1) in DMD patients **(Supplemental tables 3 and 4, respectively**), and **Figure 3a.** Among the dysregulated miRNAs, we validated the dysregulation in DMD of the dystromiRs ^24–27, 29, 32^, the heart-enriched cardiomiRs and a large number of miRNAs belonging to the DLK1-DIO3 cluster ^27^. Of the newly identified dysregulated miRNAs, we noticed in particular a large number of the Let-7 family members, the entire miR-320 family, and many miRNAs which are known modulators of diverse biological functions in skeletal and cardiac muscles, including miR-128 ^50^, miR-199a ^51, 52^, miR-223 ^53, 54^, miR-486 ^55, 56^, and members of the miR-29 ^57^ and miR-30 ^58^ families. The proximity between miR class members on the Volcano plot, suggested their similar dysregulation. Indeed, highly significant expression correlations were identified among miR members within the distinct miR classes **(Supplemental figure 2),** supporting their coordinated dysregulation. The dysregulation pattern of a selected number of miRNAs are presented graphically (**Figure 3**), including the upregulated miR-206, miR-208a-3p, miR-128-3p, miR-199a-3p and miR-369-5p **(Figure 3b)**, the downregulated miR-342-3p, miR-320a, miR-361-3p, miR-29b-3p and miR-30e-5p **(figure 3c)**, and the GC-responsive miR-223-3p, miR-379-5p, miR-27b-5p and Let-7d-5p **(Figure 3d)**. To analyze the glucocorticoid response, we compared miRNA expression before and after glucocorticoid treatment in the 4-12 age group DMD patients and identified 11 miRNAs which were dysregulated in untreated DMD patients compared to healthy controls, and for which the expression was significantly changed by the glucocorticoid treatment. Expression of nine of these miRNAs shifted toward normalization by the GC-treatment. Of note, miR-27b-5p, let-7i-5p and miR-379-5p were no longer dysregulated in the group of treated DMD versus healthy control **(Supplemental table 5)**. Similarly, miR-27b-5p and miR-379-5p were not statistically different from healthy controls when all DMD patients (treated + untreated) are grouped together.

**Figure 3.**
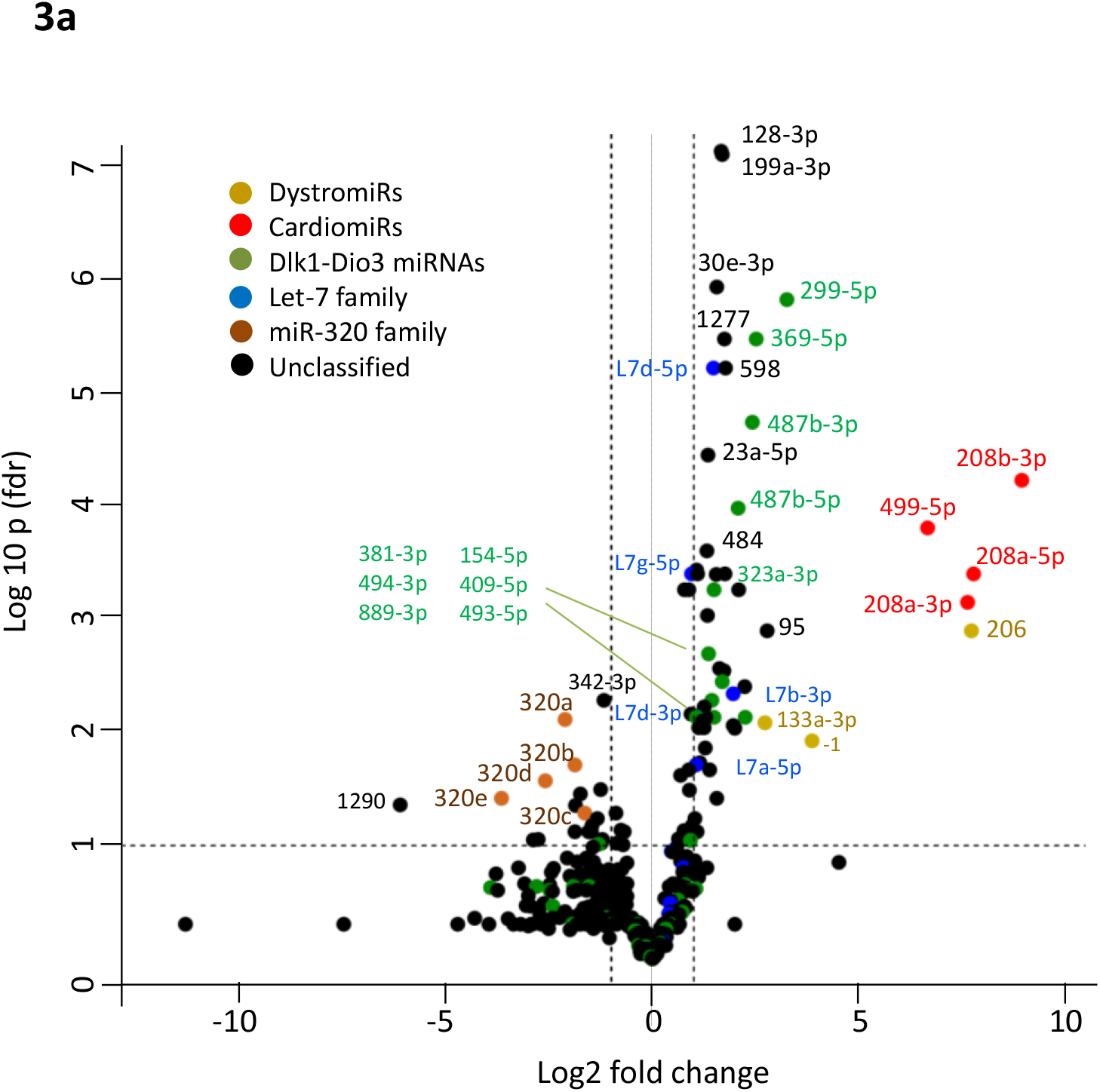
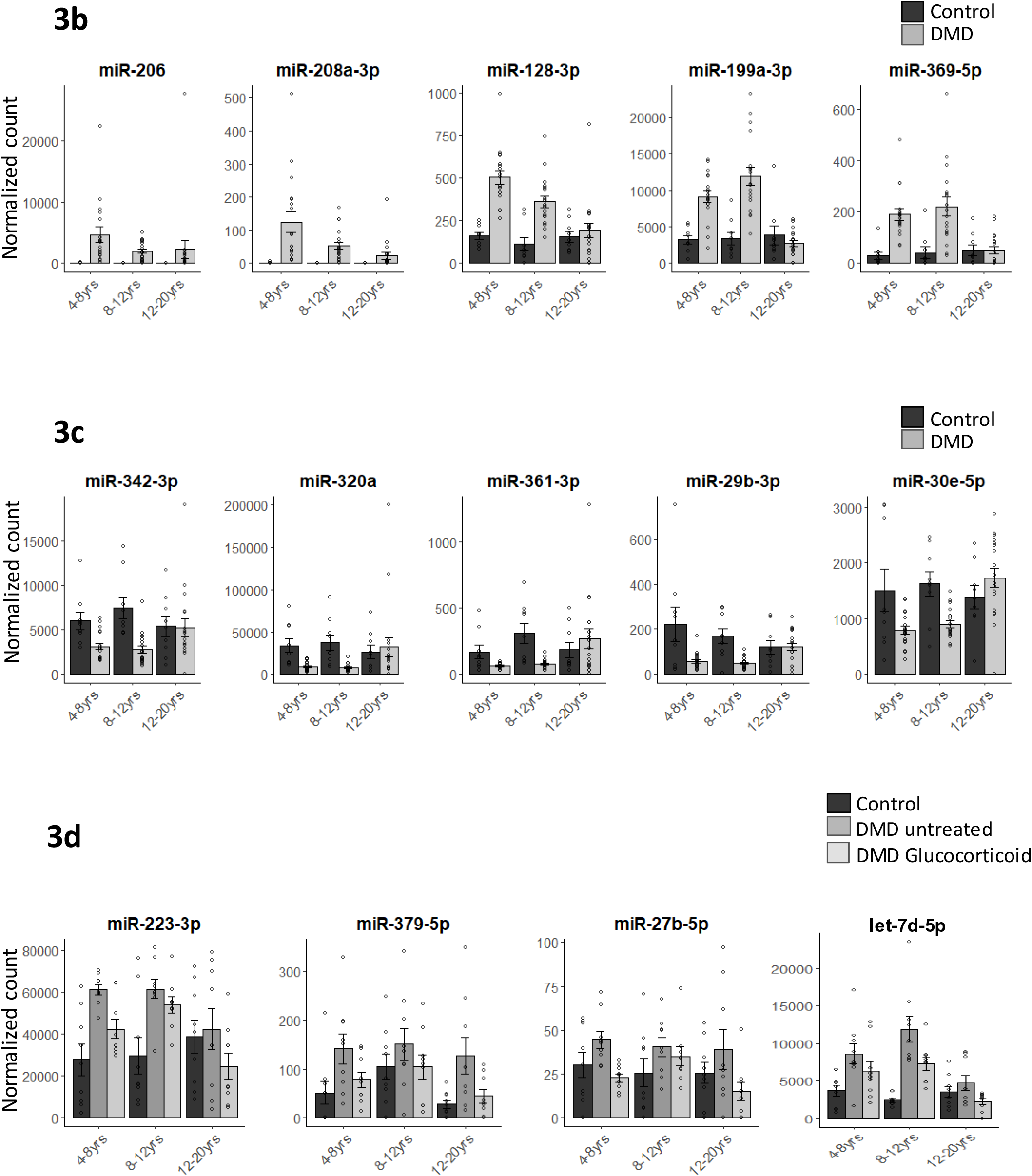
Plasma miRNA profiling in the DMD cohort. **(3a)** A volcano plot of miRNA dysregulation in the plasma of 4-12 years old DMD patients (treated & untreated together, n=36) versus healthy control (n=18). Upregulated miRNA in DMD are on the right side and downregulated on the left side of the threshold lines (FC± 1.5). MiRNAs above the horizontal line are differentially expressed with a p value <0.1 (FDR). DystromiRs in yellow, cardiomiRs in red, DLK1-DIO3 miRs in green, Let7 family miR in blue, miR-320 family in brown, unclassified miR in black. **(3b)** A graphical presentation of miRNA upregulated in DMD. **(3c)** A graphical presentation of miRNA down-regulated in DMD. **(3d)** Glucocorticoid responsive miRNAs in DMD. Each dot is one patient, n=9, error bar = SEM.

### Bioinformatics interpretation of miRNA dysregulation

*Investigation of predicted target genes for dysregulated miRNA*s: MicroRNAs are controlling the expression and/or stability of target genes, and consequently the biological functions of miRNAs are mediated by their targets. A common method for the investigation of miRNAs functions consists of the identification of signaling pathways and cellular functions of these target genes. For the identification of pathways that might be controlled by dysregulated miRNAs in DMD, we used the KEGG function of DIANA TOOLS miRPath V3 ^47^, considering all up and down miRNAs (fdr <0.1) of 4-12 years old DMD compared to age-matched controls. The analysis identified 20 distinct significantly enriched (p<0.05) pathways, including pathways already suspected of participating in the DMD disease such as Extra-cellular matrix (ECM)-receptor interaction, fatty acid biosynthesis, focal adhesion, PI3k-AKt signaling, and Foxo signaling (**Supplemental table 6)**

*Investigation of host-genes for dysregulated miRNAs in DMD*: We also attempted at the development of a complementary procedure for the biological interpretation of miRNA dysregulation. More than 1/2 of human miRNAs are embedded within introns and exons of protein coding genes ^48^. The embedded miRNAs are often co-expressed with their host genes and can regulate their expression and activity ^48, 59–62^, which support functional relations between the miRNA host-gene, target genes, and biological activity ^63, 64^. Indeed, we found that twelve of the identified dysregulated miRNAs were embedded in seven host genes that are known to be dysregulated in DMD models and patients (**Supplemental table 7**). Accordingly, we explored the possibility that analysis of host genes for dysregulated miRNAs may provide an insight into functional causes and consequences of miRNA dysregulation. The Ingenuity pathway analysis (IPA) package (Qiagen-bioinformatics) was used for the algorithmic prediction of host-gene networks, assigning the fold change and p value of their embedded miRNAs **(Supplemental table 8)**. Three networks were identified, all of which including the term Lipid Metabolism, and Small Molecule Biochemistry (**Table 1**). Of interest, the three networks integrate a number of biomolecules that are known to be involved in DMD pathophysiology, some of which are recognized as therapeutic targets, supporting the relevance of the analysis. This included, in network-1, insulin, MAPK, Erk1/2, PI3K and Akt, all members of the canonical muscle hypertrophy pathway ^65, 66^, NF-ĸB ^67–69^, H3/H4 histones, PDGF and PDGFR ^70, 71^, Hox9A ^72^, and the Mediator complex ^73^. The SREBp-1 gene was identified as the most connected in the first and largest network (**Network 1, Figure 4a)**. Network 2 includes cholesterol ^37^, EGFR ^74, 75^, the α and β estradiol and their estrogen receptor ^76^. Beta estradiol was the most connected molecule in this network (**Network 2, Figure 4b).** Network 3 includes TGF-beta ^77^ and the hepatocyte nuclear factor HNF4a, which is its most connected molecule (**Network 3, Figure 4c**). Finally, upon integration the three networks, using the IPA software, the SREBp-1 molecule was the most connected molecule of the combined network (**Figure 4d)**. This basic-helix-loop-helix transcription factor, as well as its family member SREBP2, can bind specific sterol regulatory element DNA sequences, to upregulate the synthesis of enzymes involved in sterol biosynthesis. Thus, the SREBs are master regulators of lipid metabolism ^78^.

**Figure 4.**
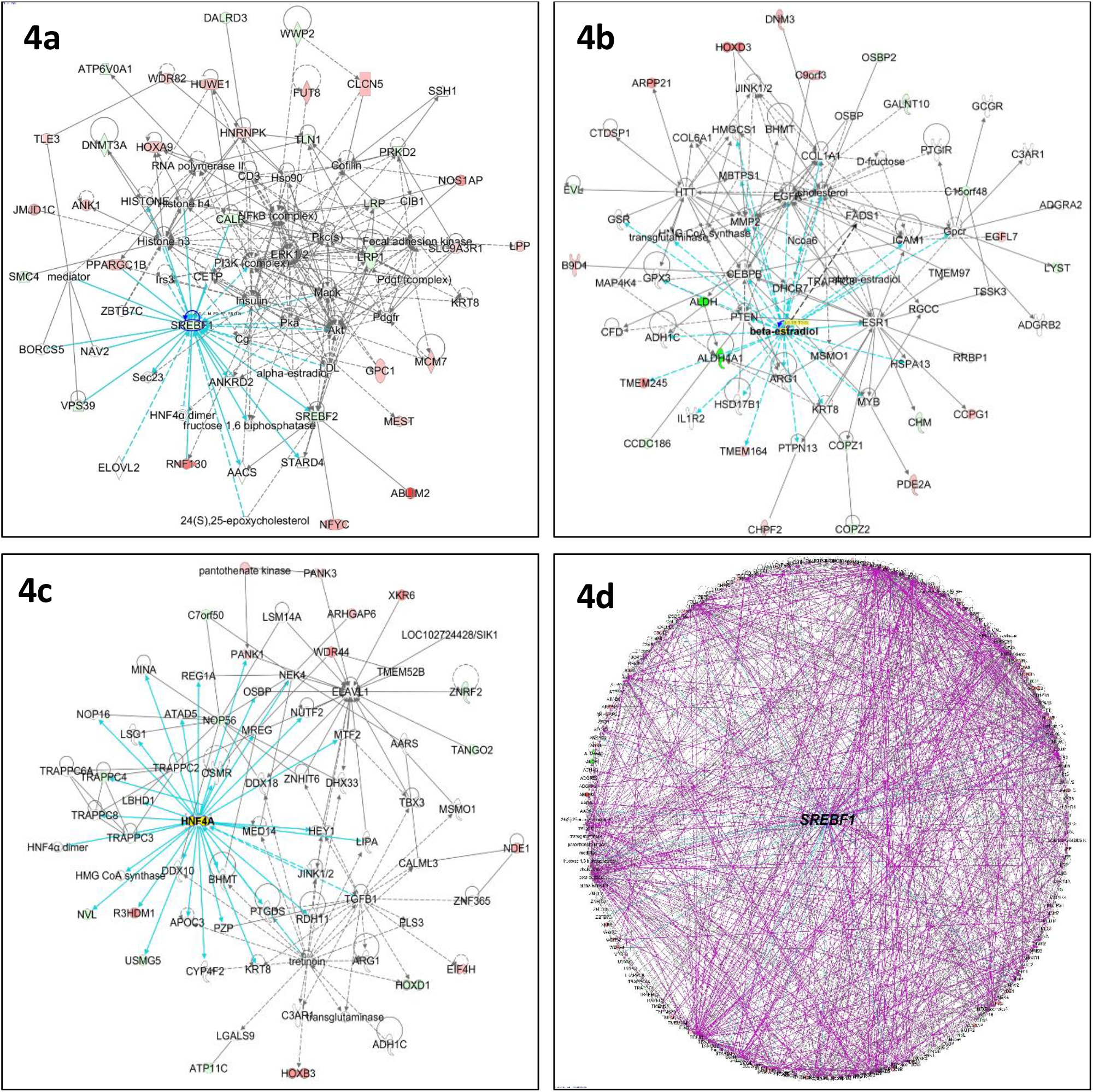
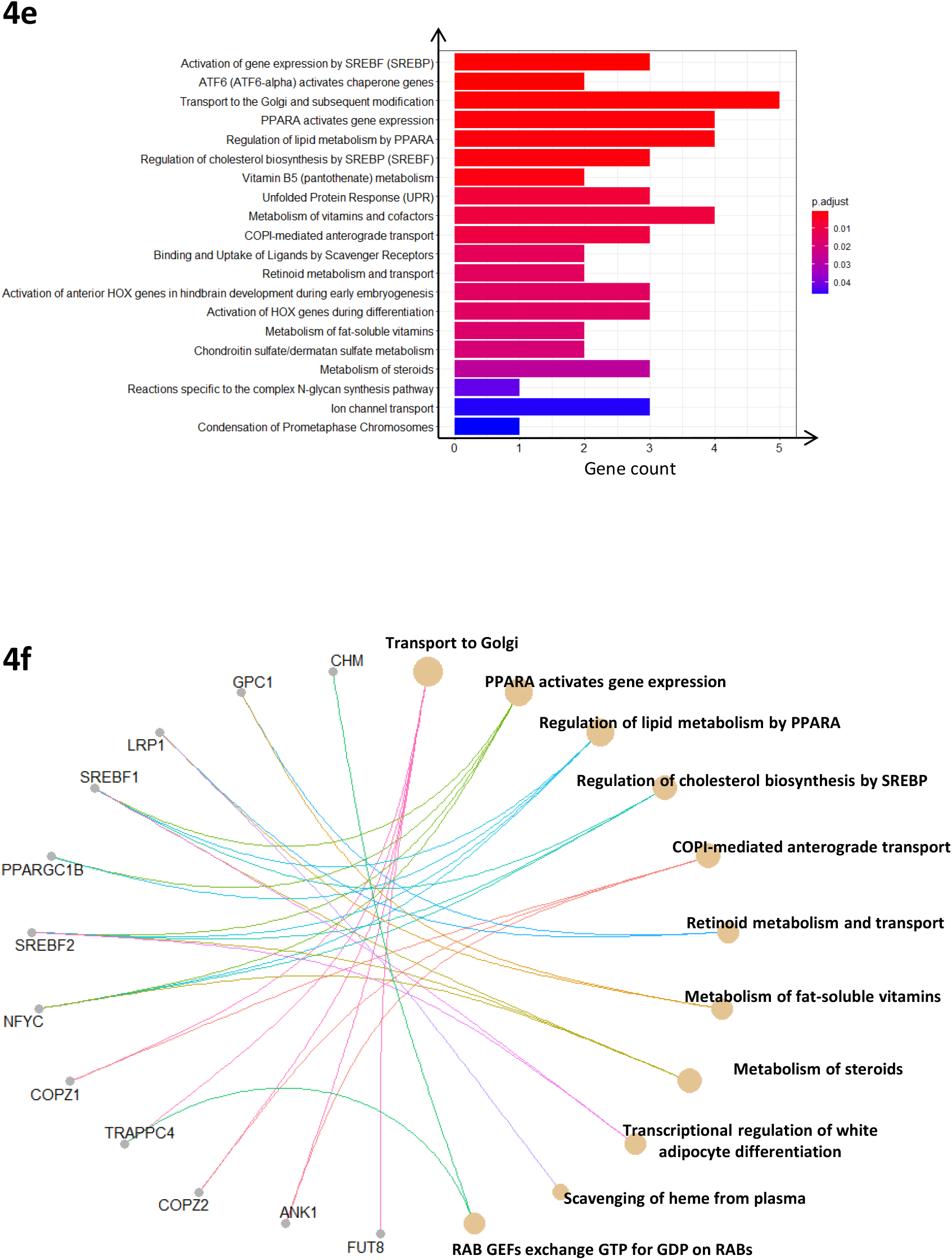
Bioinformatic analysis of host-genes for dysregulated miRNAs in DMD plasma. **(4a)** Network 1: Lipid Metabolism, Molecular Transport, Small Molecule Biochemistry, neurological disease, cancer, organismal, injury and abnormalities, **(4b)** Network 2: Lipid Metabolism, Small Molecule Biochemistry, Molecular Transport. **(4c)** Network 3, lipid metabolism, small molecule biochemistry, Dermatological Diseases and Conditions. **(4d)** Merged networks 1-3, Lipid metabolism. The connected molecules in each network are in blue. Red is upregulated, green is downregulated, continuous and discontinuous arrows are respectively direct and indirect relations, except in **4d**. Round circle = complex, rhombus = enzyme, inverse triangle = kinase, flat (horizontally oriented) circle = transcription regulator, vertically oriented circle = transmembrane receptor, trapeze = transporter. **(4e)** GO terms of host genes for dysregulated miRNAs, classified by p value. **(4f)** Lipid dysregulation network in DMD. A graphical presentation of a sub selection of miRNA host genes, which are participating in lipid metabolism, and their related GO terms.

In a complementary approach, the same 71 host genes (**Supplemental table 8**) were subjected to a gene ontology (GO) terms enrichment analysis, using the ReactomePA R algorithm. Particular enrichment was identified for terms that are related to lipid metabolism, including the activation of gene expression by SREBp (at the upper position by p value), regulation of lipid metabolism by Peroxisome Proliferator Activated Receptor Alpha (PPARA), COPI mediated transport, regulation of cholesterol biosynthesis by SREBp and metabolism of steroids **(Figure 4e)**. Considering that a large proportion of host genes are involved in lipid metabolism, the ClusterProfiler R package ^49^ was used on a sub-selection of the host genes which are known to participate in lipid metabolism. This analysis suggested the involvement of this network in the functions of transport to Golgi, PPARA transcriptional activity and regulation of lipid metabolism, regulation of cholesterol synthesis, COPI mediated anterograde transport, metabolism of fat-soluble vitamins (vitamin D) and of steroids (**Figure 4f)**. Thus, two different bioinformatics analyses identified the SERBp-dependent transcription program and the metabolism of lipid and cholesterol in the center of a network of host genes for dysregulated miRNAs in DMD.

### Evidence for lipid metabolism dysregulation in the mdx model

The miRNA host gene bioinformatics analysis predicted the dysregulation of the SREBp pathway in DMD. To investigate the relevance of this prediction, we quantified the expression of mRNA and proteins of lipid metabolism components, with a focus on the SREBp pathway **(Figure 5a)**. In addition to SREBp-1 and SREBp-2, this analysis included the SREBp upstream regulator, SCAP, ^79^, the SREBP-1 target gene, FASN, and the SREBP-2 target genes, HMGCR and LDLR, ^80, 81^. Four and three (out of the six) transcripts were significantly upregulated in gastrocnemius (GA) and diaphragm, respectively (**Figures 5b and 5c**). At the protein level, FASN, HMGCR, and SREBp-1 were upregulated in GA **(Figures 5d and 5e)** whereas the FASN, HMGCR, SCAP and SREBP1 were significantly upregulated in diaphragm **(Figures 5f and 5g)**. Thus, this analysis supported the upregulation of both SREBp-1 and SREBP-2 expressions and activities in the dystrophic muscle of the mdx mouse.

**Figure 5.**
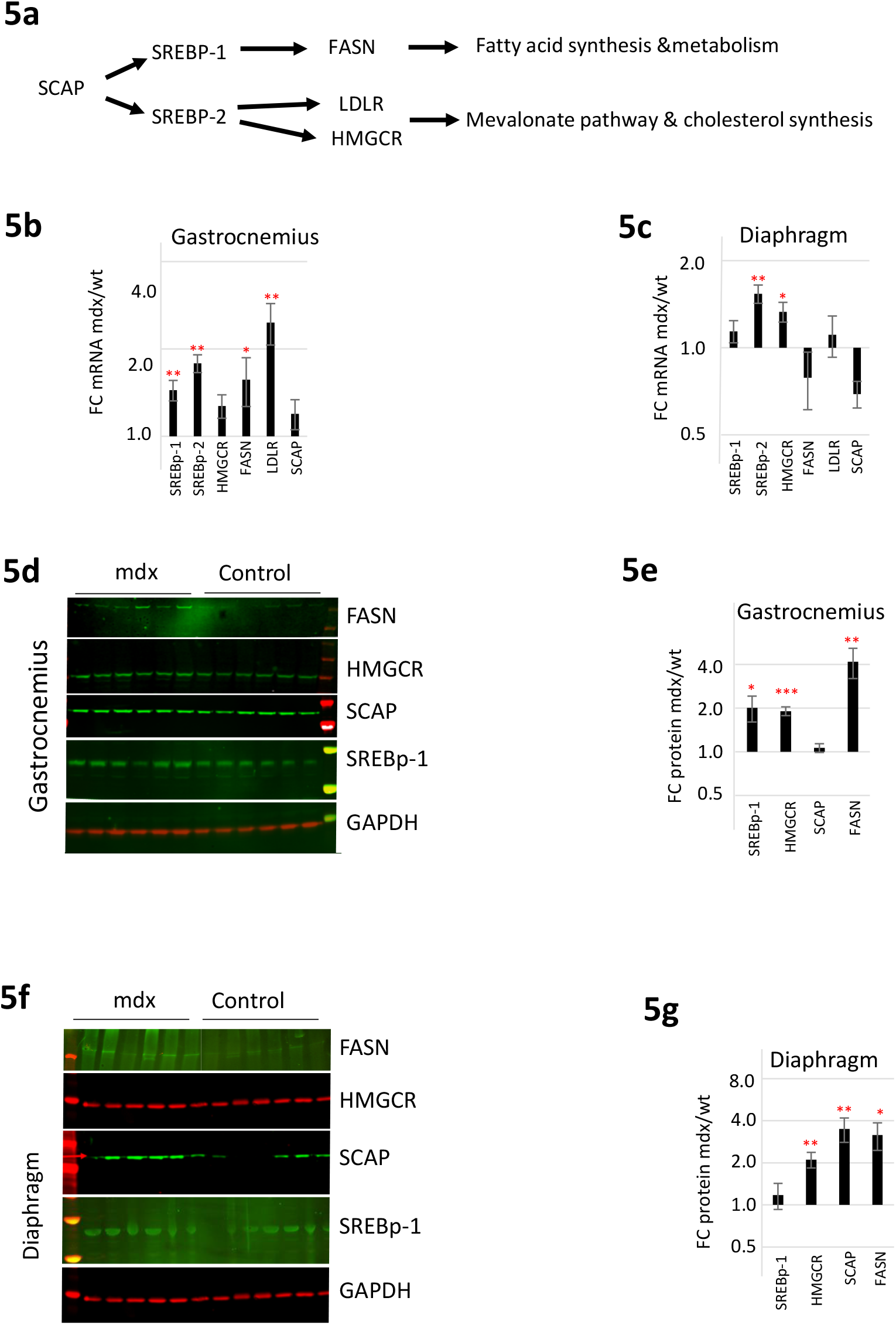
SREBp pathway in the mdx muscle. **(5a)** A schematic presentation of the components in the SREBp pathway (see text for details). **(5b-c)** Fold change SRBp pathway transcripts in the Gatsrocnemius **(5b)** and the diaphragm **(5c)** in the mdx versus healthy control mouse. **(5d-g)** A Western blot analysis of SREBP pathway protein expression in the gastrocnemius **(5d)** and its graphical quantification **(5e)**, and of the diaphragm muscle **(5f)** and its graphical quantification **(5g)**. SREBP1= Sterol regulatory binding element, SREBBP2= sterol binding regulatory element 2, HMGCR= HMG-CoA reductase, FASN= Fatty acid synthase, LDLR= Low density lipoprotein receptor, SCAP=SREBP Cleavage-Activating Protein.

### Simvastatin alleviated the dystrophic phenotype and normalized muscle cholesterol content in mdx mouse

A positive effect of the cholesterol synthesis inhibitor simvastatin on the dystrophic muscle was previously reported in the mdx mouse model ^37, 82^. The observed beneficial effect was attributed to reduction of oxidative stress, fibrosis and inflammation. Interestingly, a recent study reported an abnormally higher cholesterol content in muscles of DMD patients ^14^. Therefore, we aimed at validating the beneficial effect of simvastatin in the mdx mouse, and its possible relation with modulation of muscle cholesterol. Orally administrated simvastatin during a 3 week period significantly reduced the level of serum myomesin-3, a sensitive serum biomarker of muscular dystrophy ^83^ **(Figures 6a and 6b)**. The level of mCK was reduced as well in the simvastatin treated mdx mice (**Figure 6c**). A histological analysis of the diaphragm muscle revealed significantly reduced fibrosis in treated mdx mice **(Figure 6d and 6e**).

**Figure 6.**
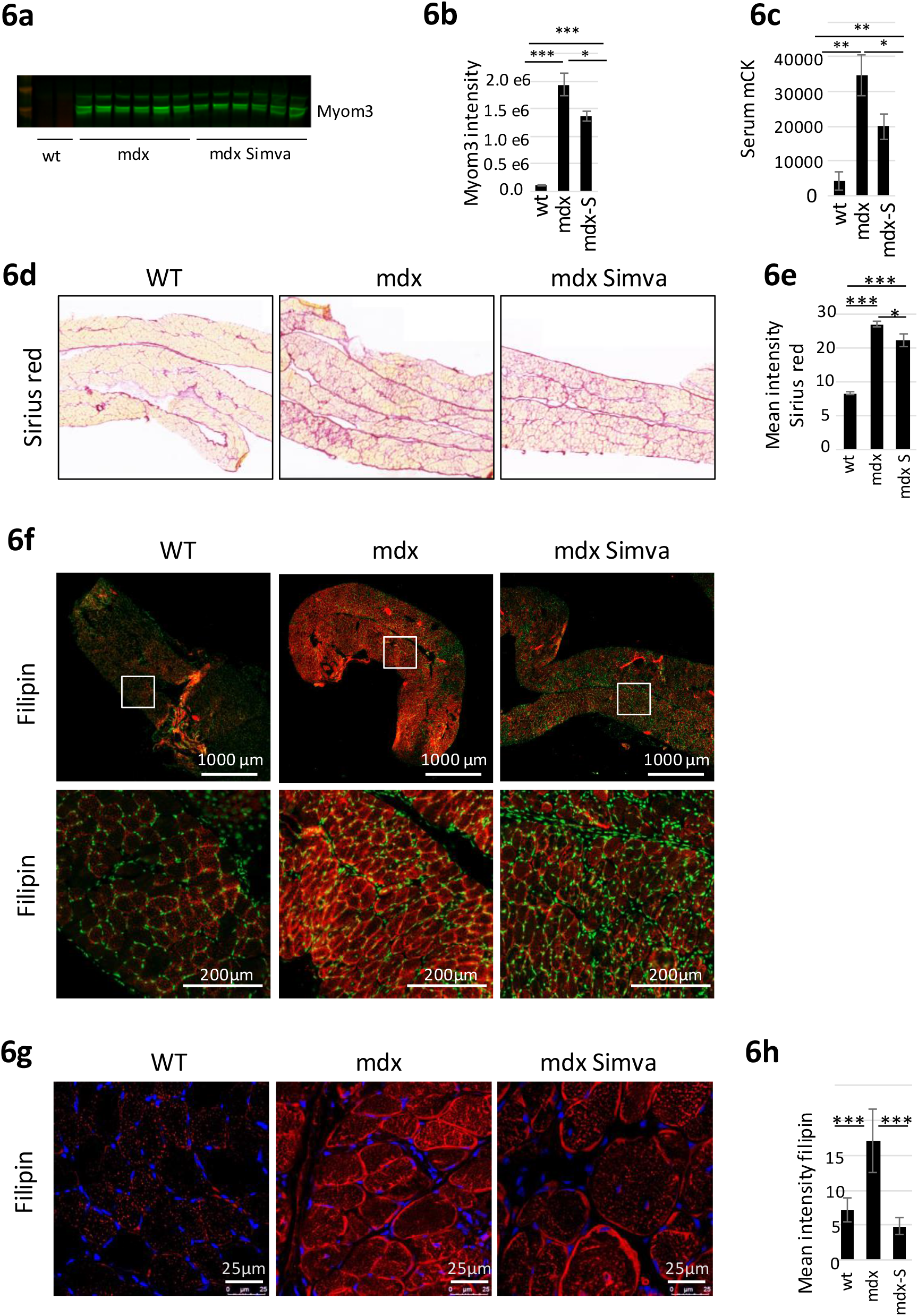
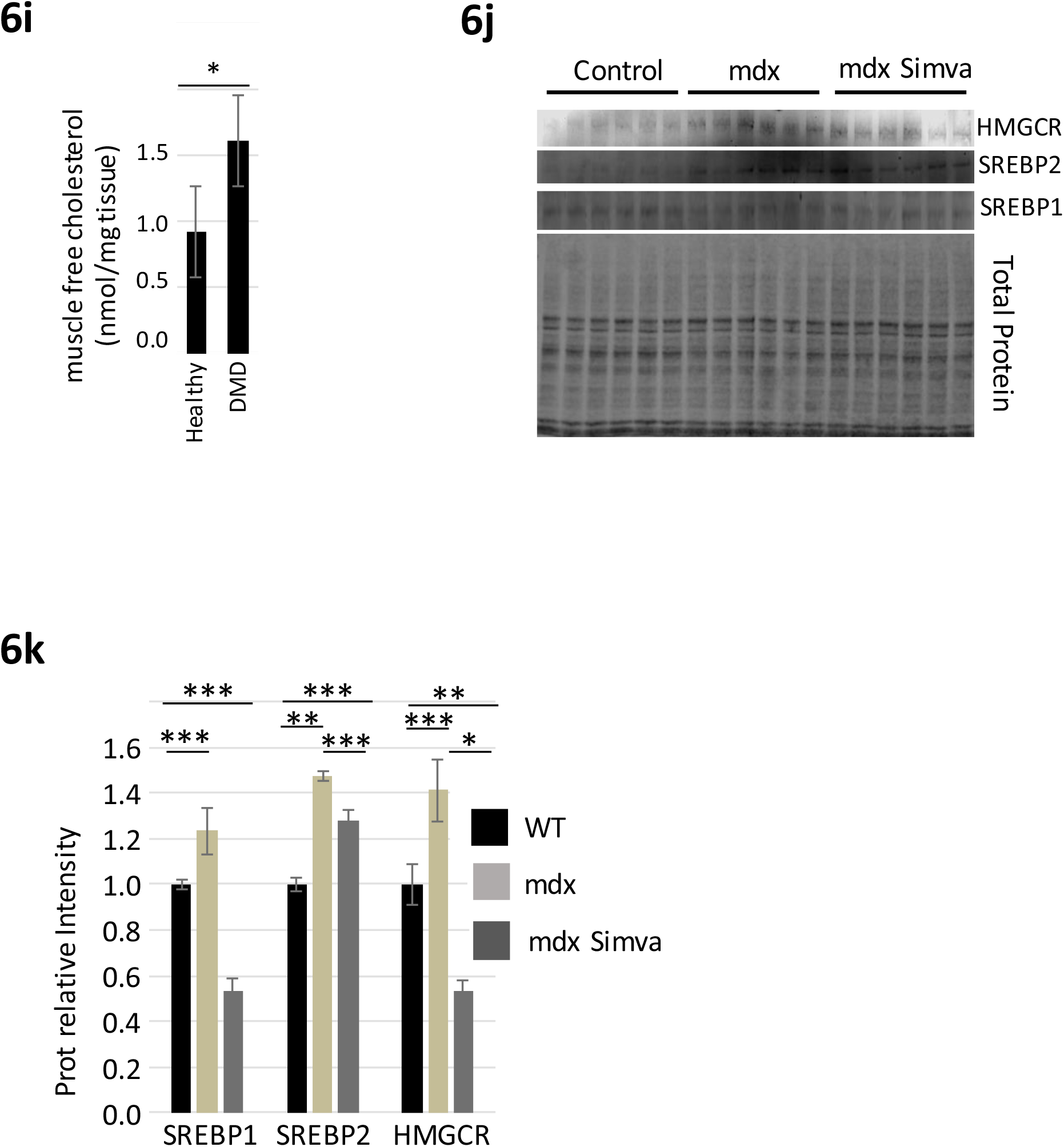
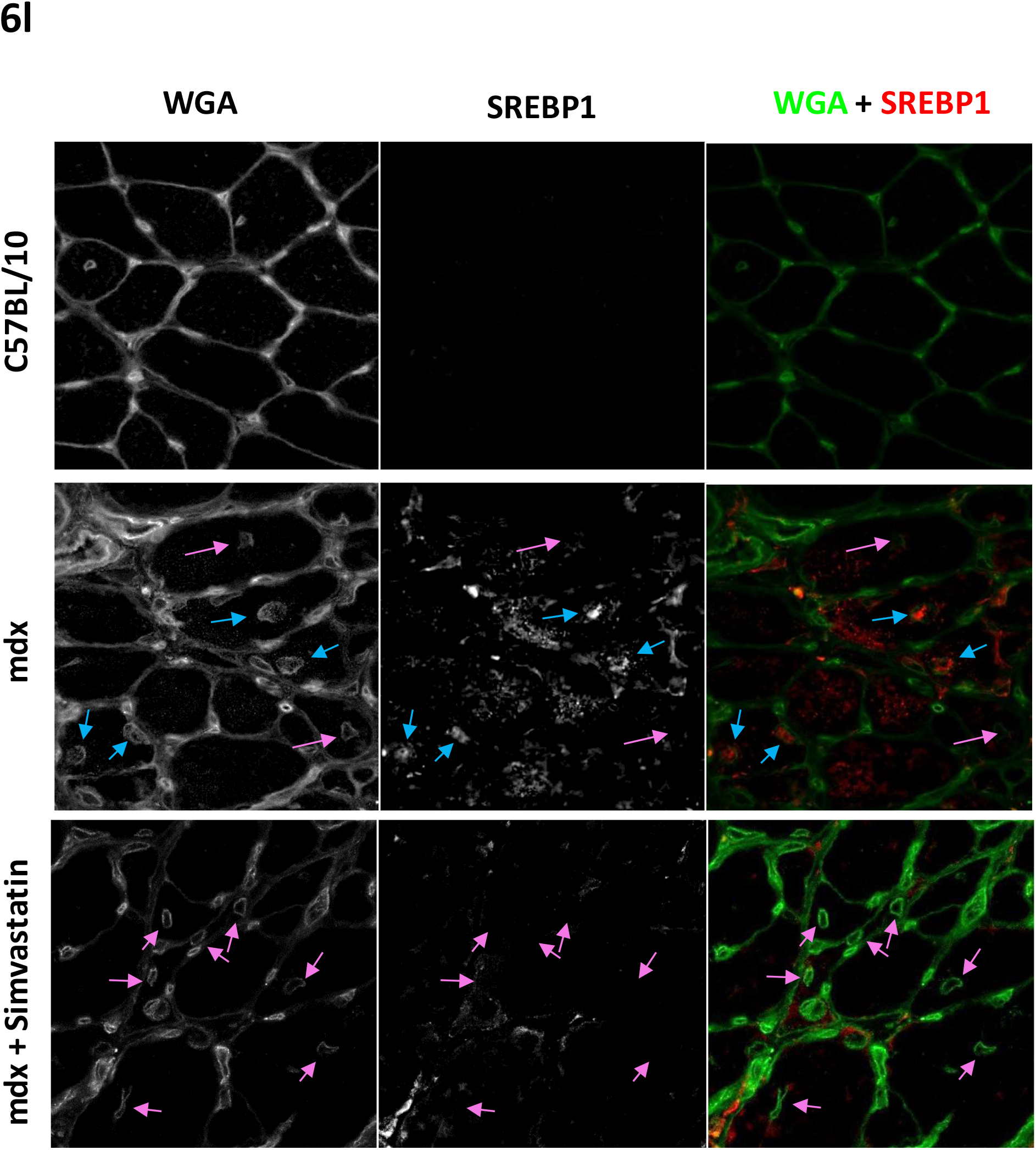
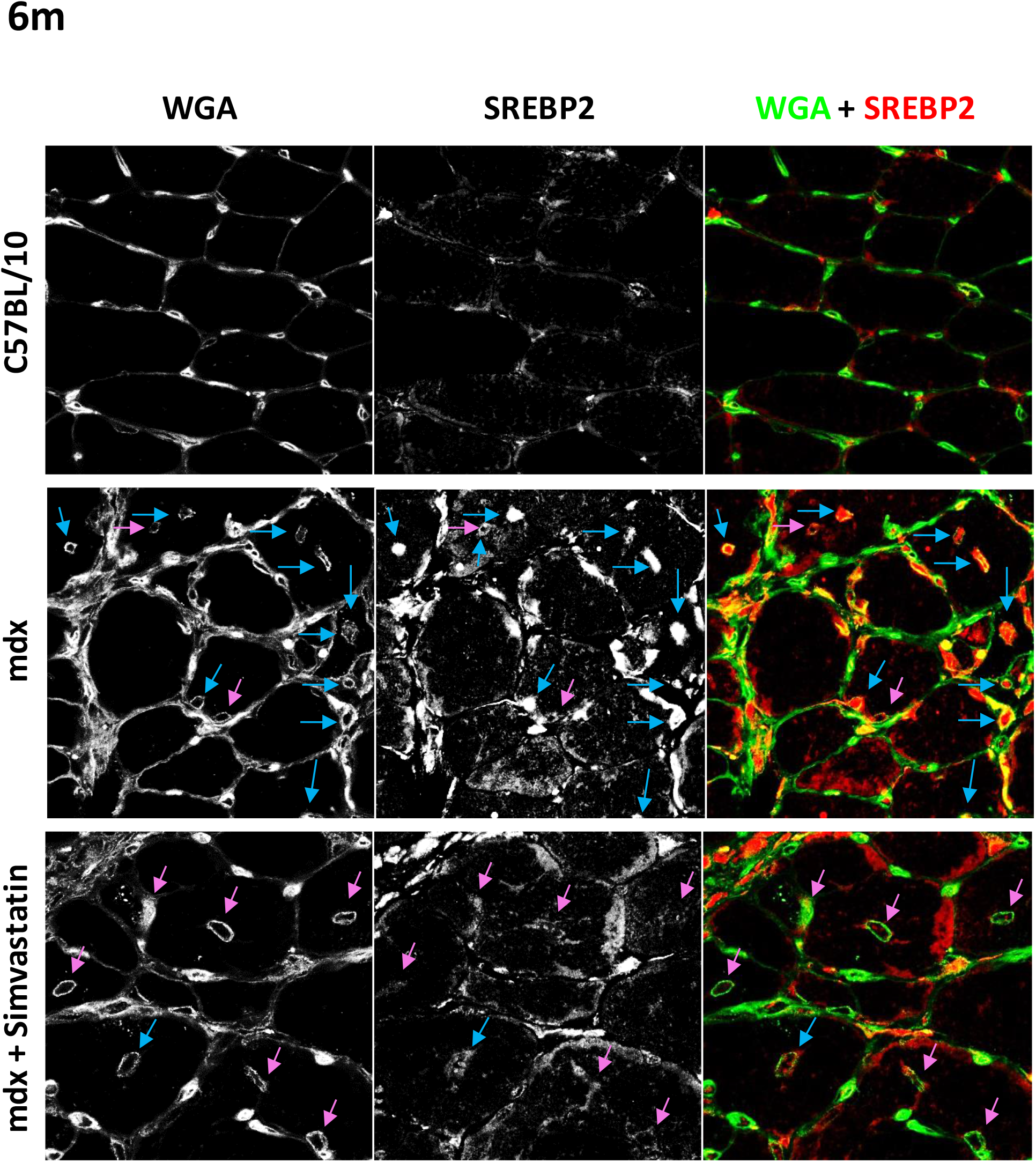
Simvastatin effect on skeletal muscle in the mdx mouse. 7-week old (young adult) control mice, and mdx mice untreated (mdx) or treated (mdx-Simva) by Simvastatin during a three weeks period (n=6)**. (6a-b)** Serum myomesin-3 (Myom3) and its graphical presentation. **(6c)** Muscle Creatine Kinase (mCK) in the blood serum. **(6d-e)** Fibrosis staining (Sirius red) of diaphragm transversal sections and its graphical presentation. **(6f)** Images of free cholesterol staining (Filipin) of transversal sections of a whole diaphragm. **(6g-h)**. Confocal images of diaphragm transversal sections **(6g)** and its quantification **(6h)**. **(6i)** Cholesterol content of skeletal muscle biopsies of DMD patients and their healthy controls. **(6j and 6k)**. A Western blot analysis **(6j)** and its graphical presentation **(6k)** of SREBP -1 SREBP-2 and HMGCR in the diaphragm muscle of control, mdx, and treated mdx mice **(6l and 6m).** Confocal microscopy images of SREBp -1, SREBp-2 in the diaphragms of the same mice. Notice that the wheat germ agglutinin (WGA) lectin stains both the sarcolemma and the myonuclear membranes. Blue arrows denote SREBP positive myonucleus inside regenerated myofibers (in the center of the myofiber). Pink arrows denote SREBP negative myonuclei in regenerated myofibers.

The level of serum cholesterol was not reduced by simvastatin **(Supplemental figure 3**), in agreement with the observation made in a previous study ^37^. We thought of investigating the consequences of simvastatin treatment at the level of the muscle tissue. Filipin is a naturally fluorescent polyene antibiotic that binds to free but not esterified sterols, and is useful therefore for the detection of free cholesterol in biological membranes ^84^. The specificity of filipin staining to cholesterol was demonstrated by treating C2C12 cells with the cholesterol-trafficking inhibitor U18666A **(Supplemental figure 4)**. Importantly, filipin staining of transversal sections confirmed increased free cholesterol content in the diaphragm of 10 weeks old mdx mice as compared to their WT controls (**Figure 6f**). Quantification of confocal images of the same sections revealed over 2-fold (P<0.001) increased cholesterol expression in the in mdx compared to wild type controls, and a complete normalization of cholesterol expression in the simvastatin treated group (p<0.001) **(Figures 6g and 6h)**. Similar reduced cholesterol content in treated mdx mice were observed also in the GA and TA muscles (**Supplemental figures 5**). Finally, the quantification of the free cholesterol content in muscle biopsies in a small cohort of human DMD patients (n=4) confirmed about two fold (p<0.035) increased free cholesterol in DMD muscles compared to controls **(Figure 6i)**. A western blot analysis demonstrated that simvastatin treatment normalized the expression of SREBP1, SREBP2 and HMGCR in the diaphragm of the mdx mouse (**Figures 6j and 6k**). Finally, we employed confocal microscopy for the investigation of SREBP-1 and SREBP-2 expression in transversal sections of the diaphragm. Higher levels of both proteins were observed in mdx, as compared to the wild type control mouse, and this expression was reversed by simvastatin **(Figures 6l and 6m).** Importantly, we noticed the expression of SRBP-1 (**Figure 6l**) and a more clearly high expression of SREBP-2 **(Figure 6m and Supplemental figure 6)** inside myonucleus in the mdx mice. The majority of the SRBP-1 and SREBP-2 positive myonucleus were in a central position in the myofibers, indicating regenerated myofibers. Of particular interest, the expression of SREBP-1 and SREBP-2 was only rarely detected in the nuclei of regenerated myofibers of the simvastatin treated mdx mice. Taken together, these results indicate that **(1)** muscles of the mdx mouse are characterized by the upregulation of the transcriptionally active nuclear forms of SREBP-1 and SREBP-2, in centrally positioned myonucleus of regenerating myofibers. **(2),** muscle fibers of the mdx mouse are characterized by increased cholesterol content, and **(3),** simvastatin treated mdx mice presented improved dystrophic phenotype in correlation with normalized muscle SREBp1 and SERBP-2 expression, and cholesterol content.

## DISCUSSION

In previous investigations, we profiled circulating miRNA in the serum of animal models for DMD ^26, 27, 34^. Following these studies, we are reporting the profiling of miRNA in the plasma of DMD patients. We validated and confirmed the dysregulation of many of the previously identified dysregulated circulating miRNAs in muscular dystrophy ^26, 27, 34^. In addition, we are reporting many newly identified dysregulated miRNA. We reasoned that an analysis of overall miRNA dysregulation can provide new information on molecular mechanisms in DMD. An original bioinformatics approach was employed, based both on target and host genes for dysregulated miRNAs. This analysis predicted a central role for the perturbation of lipid metabolism and particularly of the SREB / mevalonate / cholesterol synthesis pathway in the DMD pathology. Analysis of skeletal muscle biopsies of the mdx mouse confirmed a dysregulation of the SREBP pathway and increased cholesterol content. Treatment of mdx mice with simvastatin, an inhibitor of the mevalonate pathway and cholesterol synthesis, partially normalized the SREBP pathway, reduced the accumulation of cholesterol in the dystrophic muscles, and improved dystrophic parameters of the treated mice.

### The DMD cohort and miRNA dysregulation

To our knowledge, this is the first study to report global miRNA profiling in the plasma of DMD patients over a large age range, including GC-treated versus untreated patients. The overall miRNA dysregulation decreased in subjects above the age of 12. Thus, this age-dependent dysregulation pattern, which was shown previously with the mCK and the myomiRs, was extended in the present study to a large majority of the dysregulated miRNAs in DMD. This pattern may reflect both the reduced muscle mass and physical activity of the older DMD age group (quantitative changes), as well pathophysiological evolution, independently of muscle mass and activity (qualitative changes). We confirmed the upregulation of many miRNAs that were identified in previous investigations. These include the dystromiRs, the cardiomiRs, and the DLK1-DIO3 clustered miRNAs ^26, 27, 85^. Of interest, we are reporting here, to our knowledge for the first time in DMD, the dysregulation of many members of the Let-7 and miR-320 families. Among the newly identified dysregulated miRNA in DMD plasma, we identified many miRNAs that are known to play diverse roles in muscle pathophysiology. Among these are miR-128 ^50, 86^, miR-199a ^51^, miR-223 ^53^, miR-486 ^55, 56^, and the miR-29 ^87^ and miR-30 ^58, 88^ family members. As in previous investigations, we identified a higher number of upregulated rather than downregulated circulating miRNA in DMD. It is thought the upregulation in DMD of circulating miRNAs results principally from increased passive leakiness from damaged fibers and the active passage of miRNAs through the dystrophic sarcolemma. Of interest, however, in the present study, is the large number of downregulated miRNAs which we identified in the plasma in DMD. The downregulation of circulating miRNA is not explained by the alteration of sarcolemma permeability. Thus, it is likely to result from transcriptional adaptation in dystrophic tissues. Of the downregulated circulating miRNAs, we noticed in particular miR-342 and miR-185, both of which target the SREBp mevalonate pathway and the synthesis of cholesterol ^89–91^. Additionally, we noted the downregulation of the entire (5 members) miR-320 family, of which the biological consequences in the context of muscular dystrophy are as yet unknown.

### A holistic bioinformatics analysis of miRNA dysregulation

The conventional method for the biological interpretation of miRNA dysregulation is based on the analysis of the consequences of miRNAs dysregulation. This interpretation can be carried out by a pathway enrichment analysis of mRNA targets for the dysregulated miRNAs. The miRNA target genes analysis predicted enrichment for the KEGG pathways, of the proteoglycan ECM-receptor interaction ^92, 93^, focal adhesion ^94^, fatty acid biosynthesis ^13, 14^, as well as several signaling pathways : Hippo ^95^, PI3K-Akt ^96^, estrogen ^97^, FoxO ^98^ and ErbB ^99^, all of which are known to be dysregulated in DMD. The identification of the dysregulation of these pathway is well-known in the field and thus supports the pertinence of our miRNA dysregulation results.

However, the target genes analysis failed to produce novel hypotheses or an explanation for the DMD pathophysiology. Importantly, among the dysregulated miRNAs, we noticed some which are positioned inside mRNA transcripts that are known to be dysregulated in the dystrophic muscle in DMD. This observation supports a hypothesis that, in DMD, miRNAs and their host-gene might be coordinately dysregulated, and thus, that miRNA dysregulation may indicate a host-gene dysregulation. As mentioned above, in the analysis which is based on miRNAs target genes, the focus is on the consequences of the dysregulation of the miRNAs. In contrast, in the host gene approach, the focus is on the causes of the dysregulation of the miRNAs, i.e. on the upstream events of miRNA dysregulation, thus providing different information. The host gene approach is of particular interest for the interpretation of miRNA dysregulation in the serum/plasma, since mRNA profiling cannot be carried out in this compartment (that express principally remnant of degraded mRNAs). Taking into account all these considerations, we hypothesized that a biological interpretation approach for circulating miRNA that combine both target and host genes (described schematically in **supplemental Figure 7**), may provide an improved interpretation of DMD pathophysiology. To predict the possible consequences of dysregulation of the host genes, we used two different approaches. In the first, we used the IPA algorithm for the construction of gene networks, and in the second, we used the ReactomePA and ClusterProfiler R algorithms for the identification and illustration of GO terms, which are associated with the host genes. The IPA-based gene-interaction analysis predicted three dysregulated networks, all of which contain the term lipid metabolism. At the central positions of the three networks, which identify the most connected molecules, we found SREBP1, Beta estradiol, and HNFA4. The SREBp 1 molecule was identified as being central to the merged network of dysregulation in DMD. Similarly, the analysis of GO terms by the ReactomPA algorithm predicted a key role for the SREBp transcription factors. Importantly, the GO term analysis supports the hypothesis that the dysregulation of the SREBp pathway may happen in the context of a broader level lipid metabolism dysregulation as observed with the Reactome IPA analysis.

### Lipid metabolism dysregulation in the mdx mouse

The SREBP family of transcription factors is composed of three members. The two isoforms SREBP1c and SREBP1a, result from two distinct promoters of the SREBPF1 gene. SREBP-1 isoforms control primarily lipogenic gene expression, while SREBP-2 regulates the transcription of genes related to cholesterol metabolism ^81^. Indeed, in agreement with the bioinformatics prediction, gene expression analysis confirmed the significant upregulation of members of the SREBp pathway in the gastrocnemius and the diaphragm muscles in the young mdx mouse, providing supporting experimental data from the dystrophic muscle for the hypothesis of SREB pathway dysregulation.

### Treating mice by simvastatin

The mevalonate pathway and cholesterol synthesis are inhibited by statins, by acting directly on the HMGCR, the rate limiting enzyme of the cholesterol synthesis pathway. The treatment of mdx mice with the FDA-approved drug simvastatin was reported to affect positively the mdx mice, which presented reduced dystrophic symptoms and improved muscle function. Simvastatin is a pleotropic drug and its positive effect on mdx was reported to be independent of lipid metabolism in accordance with the absence of modification of the seric cholesterol level ^37^. Importantly, we identified an overexpressed mevalonate pathway in the skeletal muscle of the mdx mouse, as well as increased cholesterol content, both of which returned toward normal after simvastatin treatment. Thus, while the positive effect of simvastatin was repeatedly suggested, the molecular basis for this effect was not clear ^37, 82^. In the present investigation, we identified a positive correlation between the effect of simvastatin on the muscle mevalonate pathway, and its alleviated dystrophic parameters in the mdx mouse. This correlation supports that HMGCR and accumulation of muscle cholesterol are the therapeutic targets of simvastatin in the dystrophic muscle. The molecular mechanism by which the increased cholesterol level affecting the dystrophic muscle in DMD is yet unknown.

However, our investigation is in agreement with earlier reports in support of lipids and particularly of cholesterol metabolism abnormalities in DMD ^13,14,15, 100–105, 106–112^. Of particular interest, Steen and colleagues reported the improved dystrophic parameters in an mdx mouse that overexpress the NPC1 gene, an accelerator of intracellular cholesterol trafficking ^113^, while Milad and colleagues reported the correlation between increased plasma cholesterol and accentuated muscular dystrophy in the mdx mouse ^112^, which together supporting that perturbation of cholesterol metabolism affects the DMD phenotype. These early reports failed, however, to provide a molecular hypothesis for the causes and origin of this dysregulation. Not only does our study concords with these previous findings, it also provides a molecular framework for this dysregulation for the first time. In summary, from the plasma samples of a large DMD cohort, the present study identified that perturbation of lipid metabolism, and in particular of the cholesterol homeostasis, play an important role in the pathophysiology of DMD. These findings are opening novel perspective for clinical interventions in DMD.

## AUTHOR CONTRIBUTION

**FA** and **AVH** designed and performed experiments, analyzed results and wrote the paper. **MS** and **LSu** performed experiments and analyzed results. **GC** performed bioinformatics analyses. **LSe** and **TV** designed experiments, managed the clinical trial and analyzed results. **SB** coordinated the clinical trial. **IR** and **DI** designed experiments, managed the project and wrote the manuscript.

## ACKNOWLEDGEMENT

We are grateful to the *In vivo* evaluation, Imaging Histology services of Genethon and to Siân Cronin for critical reading of the manuscript. This study was financially supported by the Association Francais contre les Myopathies (AFM); by ADNA (Advanced Diagnostics for New Therapeutic Approaches); the Institut National de la Sante et de la Recherche Medicale (INSERM); the Universite Pierre et Marie Curie Paris 06; the Centre National de la Recherche Scientifique (CNRS). The authors wish to thank patients and parents for participating, and all staff involved in sample collections in the different medical centers. The authors of this manuscript certify that they comply with the ethical guidelines for authorship and publishing in the Journal of Cachexia, Sarcopenia and Muscle.

## SUPPLEMENTAL DATA

**Supplemental table 1.** Cohort’s medical center composition

|  | **City** | **medical center** | **DMD** | **Control** | **principal investigator** |
| --- | --- | --- | --- | --- | --- |
| 1 | Liege | CHR Citadelle | 22 | 24 | Dr Laurent SERVAIS |
| 2 | Gent | UZ Ghent | 13 | 4 | Dr Nicolas DECONINCK |
| 3 | Bucarest | Al Obregia Hospital | 16 | 45 | Dr Dana CRAIU |
| 4 | Paris | AIM | 22 | 17 | Dr Laurent SERVAIS |
| 5 | Amiens | Hôpital Nord | 3 | 9 | Dr Anne Gaëlle LE MOING |
| 6 | Rennes | Hôpital Pontchaillou | 2 | 0 | Dr Hélène RAUSCENT |
| 7 | Lille | Hôpital Roger Salengro | 3 | 18 | Dr Jean Marie CUISSET |
| 8 | Toulouse | Hôpital des Enfants CHU Purpan | 10 | 0 | Dr Claude CANCES |
| 9 | Nantes | CHU Hôtel Dieu | 7 | 5 | Pr Yann PEREON |
| 10 | Garches | Hôpital Raymond-Poincaré | 2 | 1 | Dr Brigitte ESTOURNET |
| Total |  |  | 100 | 123 |  |

**Supplemental table 2.** Clinical features of the DMD patients. Glucocorticoid treated (T) patients are in blue. Untreated patients (UT) are in black. (*) from Martinique; **(**)** Mother from the Reunion island and Martinique, Father Caucasian, (***) Mother from Brazil, Father Caucasian. NA: not available. ND: not done.

| HTS ID | Group | Country | Ethnic origin | Age (Years) | Weight (kg) | Height (cm) | Gene mutation confirming diagnosis | Age of onset of first symptoms (years) | Loss of ability to walk | Age ability to walk lost (years) | Walton scale | Brooke scale | CPK | Cardiac ejection fraction (EF) |
| --- | --- | --- | --- | --- | --- | --- | --- | --- | --- | --- | --- | --- | --- | --- |
| D1 | 4-8 UT | France | Caucasian | 4 | 16.8 | 98 | Del exons 53 - 54 | 3 | No | . | 1 | 1 | 9860 | 65 |
| D2 | 4-8 UT | Belgium | Caucasian | 4 | 16.9 | 102 | Del exons 46 - 48 | 2 | No | . | 1 | 1 | 22158 | ND |
| D3 | 4-8 UT | France | Asian | 4 | 19.1 | 107 | Del exon 51 | 3 | No | . | 3 | 1 | 20391 | ND |
| D4 | 4-8 UT | France | Caucasian | 4 | 16.9 | 108 | Del T in exon 74 | 2 | No | . | 2 | 1 | 16067 | 65 |
| D5 | 4-8 UT | France | Caucasian | 4 | 17.2 | 107 | Del exons 46 - 48 | 3 | No | . | 2 | 1 | 18408 | ND |
| D6 | 4-8 UT | France | Caucasian | 6 | 23.8 | 119 | Point mutation exon 23 | 2 | No | . | 3 | 1 | 1392 | ND |
| D7 | 4-8 UT | Belgium | African | 7 | 18 | 110 | Splicing mutation exon 67 | <1 | No | . | 3 | 1 | 5597 | 65 |
| D8 | 4-8 UT | Belgium | Caucasian | 7 | 23.2 | 116 | Del exons 48 - 54 | 3 | No | . | 1 | 1 | 14002 | ND |
| D9 | 4-8 UT | Belgium | Caucasian | 7 | 21.5 | 122 | Del exons 3 - 7 | NA | No | . | 3 | 1 | 5997 | 67 |
| D10 | 4-8 T | Belgium | Caucasian | 4 | 17.2 | 105 | Del exons 46 - 48 | 2 | No | . | 1 | 1 | 14157 | 71 |
| D11 | 4-8 T | France | Caucasian | 4 | 17.5 | 101 | Del exons 53 - 54 | 3 | No | . | 4 | 1 | 17808 | ND |
| D12 | 4-8 T | France | Asian | 4 | 20.5 | 109 | Del exon 51 | 3 | No | . |  | 1 | 18924 | ND |
| D13 | 4-8 T | France | Caucasian | 4 | 17.3 | 109 | Del exons 46 - 48 | 3 | No | . | 3 | 1 | 17077 | ND |
| D14 | 4-8 T | France | Caucasian | 4 | 18.7 | 111 | Del T in exon 74 | 2 | No | . | 1 | 1 | 29539 | ND |
| D15 | 4-8 T | France | Caucasian | 6 | 26.9 | 122 | Point mutation exon 23 | 2 | No | . | 1 | 1 | 17334 | ND |
| D16 | 4-8 T | Belgium | African | 7 | 19 | 115 | Splicing mutation exon 67 | <1 | No | . | 3 | 1 | 5877 | ND |
| D17 | 4-8 T | Belgium | Caucasian | 5 | 16.8 | 108 | Del exons 51 - 55 | 3 | No | . | 1 | 1 | 23349 | ND |
| D18 | 4-8 T | Belgium | Caucasian | 7 | 23 | 122 | Del exons 3 - 7 | NA | No | . | 1 | 1 | 7887 | 65 |
| D19 | 8-12 UT | France | Caucasian | 8 | 23 | 116 | Del exon 50 | 1 | No | . | 3 | 1 | 14115 | ND |
| D20 | 8-12 UT | France | Caucasian | 8 | 35.6 | 134 | Dup exon 17 | 2 | Yes | 8 | 7 | 2 | 5763 | ND |
| D21 | 8-12 UT | France | Caucasian | 8 | 34 | 116 | Stop exon 26 | 2 | Yes | 7 | 6 | 5 | 3031 | ND |
| D22 | 8-12 UT | Romania | Caucasian | 8 | 30 | 132 | Del exon 45 | 4 | No | . | 3 | 1 | 15324 | ND |
| D23 | 8-12 UT | Belgium | Caucasian | 9 | 24 | 127 | Del 4 bp in exon 62 | 3 | No | . | 3 | 1 | 12055 | 65 |
| D24 | 8-12 UT | France | Caucasian | 9 | 26 | 122 | Stop exon 2 | 4 | No | . | 4 | 1 | 14211 | 68 |
| D25 | 8-12 UT | Romania | Caucasian | 10 | 47 | 140 | Del exon 45 | 3 | No | . | 3 | 2 | 10407 | 60 |
| D26 | 8-12 UT | France | African | 10 | 27 | 128 | Stop exon 34 | 4 | No | . | 3 | 1 | 12544 | 63 |
| D27 | 8-12 UT | France | Caucasian | 10 | 30 | 132 | Dup exon 52 - 54 | 3 | Yes | 9 | 6 | 6 | 3796 | ND |
| D28 | 8-12 T | France | Caucasian | 8 | 25.4 | 119 | Dup exon 13 - 17 | 6 | No | . | 3 | 1 | 12800 | 64 |
| D29 | 8-12 T | France | Caucasian | 8 | 24.6 | 120 | Del exons 48-50 | 2 | No | . | 3 | 1 | 24120 | ND |
| D30 | 8-12 T | France | Mixed (*) | 8 | 29 | 129 | Del 5 bp in exon 40 | 5 | No | . | 3 | 1 | 11729 | ND |
| D31 | 8-12 T | France | Caucasian | 9 | 27 | 127 | Del 22 bp in exon 68 | 1 | No | . | 3 | 1 | NA | ND |
| D32 | 8-12 T | Romania | Caucasian | 9 | 21 | 115 | Del exons 8 - 9 | 4 | No | . | 4 | 1 | 9420 | 60 |
| D33 | 8-12 T | Romania | Caucasian | 10 | 23 | 124 | Dup exons 5 - 6 | 2 | No | . | 4 | 1 | 9950 | 65 |
| D34 | 8-12 T | Romania | Caucasian | 10 | 45 | 136 | Del exon 50 | 6 | No | . | 4 | 2 | 5985 | ND |
| D35 | 8-12 T | Belgium | Caucasian | 11 | 36.5 | 134 | Dup exons 3 - 9 | 3 | No | . | 0 | 1 | 7001 | ND |
| D36 | 8-12 T | France | Mixed (**) | 12 | 27.9 | 139 | Dup exons 18 - 44 | 3 | No | . | 4 | 1 | NA | 66 |
| D37 | 12-20 UT | Belgium | Caucasian | 12 | 48 | 156 | Del exon 65 | NA | Yes | 10 | 7 | 6 | 1687 | 68 |
| D38 | 12-20 UT | Belgium | Caucasian | 13 | 50 | 162 | Del exons 48 - 50 | 4 | Yes | 10.5 | 7 | 4 | 1601 | 66 |
| D39 | 12-20 UT | France | Caucasian | 14 | 29 | 146 | Del exon 30 | 4 | Yes | 10 | 5 | 5 | 3660 | ND |
| D40 | 12-20 UT | Romania | Caucasian | 15 | 56 | 150 | Del exons 3 - 42 | 3 | Yes | 9 | 5 | 5 | 3210 | 50 |
| D41 | 12-20 UT | France | Caucasian | 16 | 52 | 175 | Del exons 45 - 50 | 2 | Yes | 8 | 5 | 5 | 658 | 40 |
| D42 | 12-20 UT | Belgium | Caucasian | 17 | 70 | 152 | Del exons 5 - 7 | 1 | Yes | 10 | 6 | 5 | 1391 | A |
| D43 | 12-20 UT | Belgium | Caucasian | 18 | 63 | 165 | Dup exons 56 - 63 | 3 | Yes | 10 | 6 | 5 | 688 | ND |
| D44 | 12-20 UT | France | Caucasian | 19 | 61 | 166 | Del exons 45- 52 | 1 | Yes | 7 | 6 | 5 | 1435 | ND |
| D45 | 12-20 UT | Belgium | Caucasian | 20 | 85 | 165 | Del exon 22 | 4 | Yes | 9 | 5 | 5 | 1779 | 45 |
| D46 | 12-20 T | Romania | Caucasian | 12 | 38 | 146 | Del exons 8 - 12 | 6 | Yes | 10 | 7 | 6 | 4138 | 65 |
| D47 | 12-20 T | France | Mixed (***) | 13 | 45 | 165 | Dup exons 3 - 42 | 3 | Yes | 10 | 7 | 6 | NA | 63 |
| D48 | 12-20 T | Belgium | Caucasian | 13 | 60.8 | 170 | Dup exons 8 - 11 | 3 | Yes | 9 | 7 | 6 | 1945 | ND |
| D49 | 12-20 T | Belgium | Caucasian | 13 | 46.5 | 139 | Del exons 46 - 52 | 3 | Yes | 12 | 4 | 1 | 6902 | ND |
| D50 | 12-20 T | Romania | Caucasian | 13 | 45 | 133 | Del exons 50 - 52 | 5 | No | . | 4 | 6 | 1254 | 60 |
| D51 | 12-20 T | Romania | Caucasian | 13 | 34 | 132 | Del exons 45 - 50 | 5 | Yes | 12 | 7 | 6 | 7547 | ND |
| D52 | 12-20 T | Romania | Caucasian | 14 | 42.5 | 143 | Del exons 45 - 50 | 6 | No | . | 3 | 1 | 4249 | ND |
| D53 | 12-20 T | France | Caucasian | 14 | 55.5 | 147 | Dup G in exon 53 | 4 | Yes | 13 | 7 | 1 | NA | 55 |
| D54 | 12-20 T | Belgium | Caucasian | 17 | 49 | 160 | Del exons 46 - 47 | 4 | Yes | 10 | 6 | 4 | 1159 | ND |

**Supplemental table 3.**
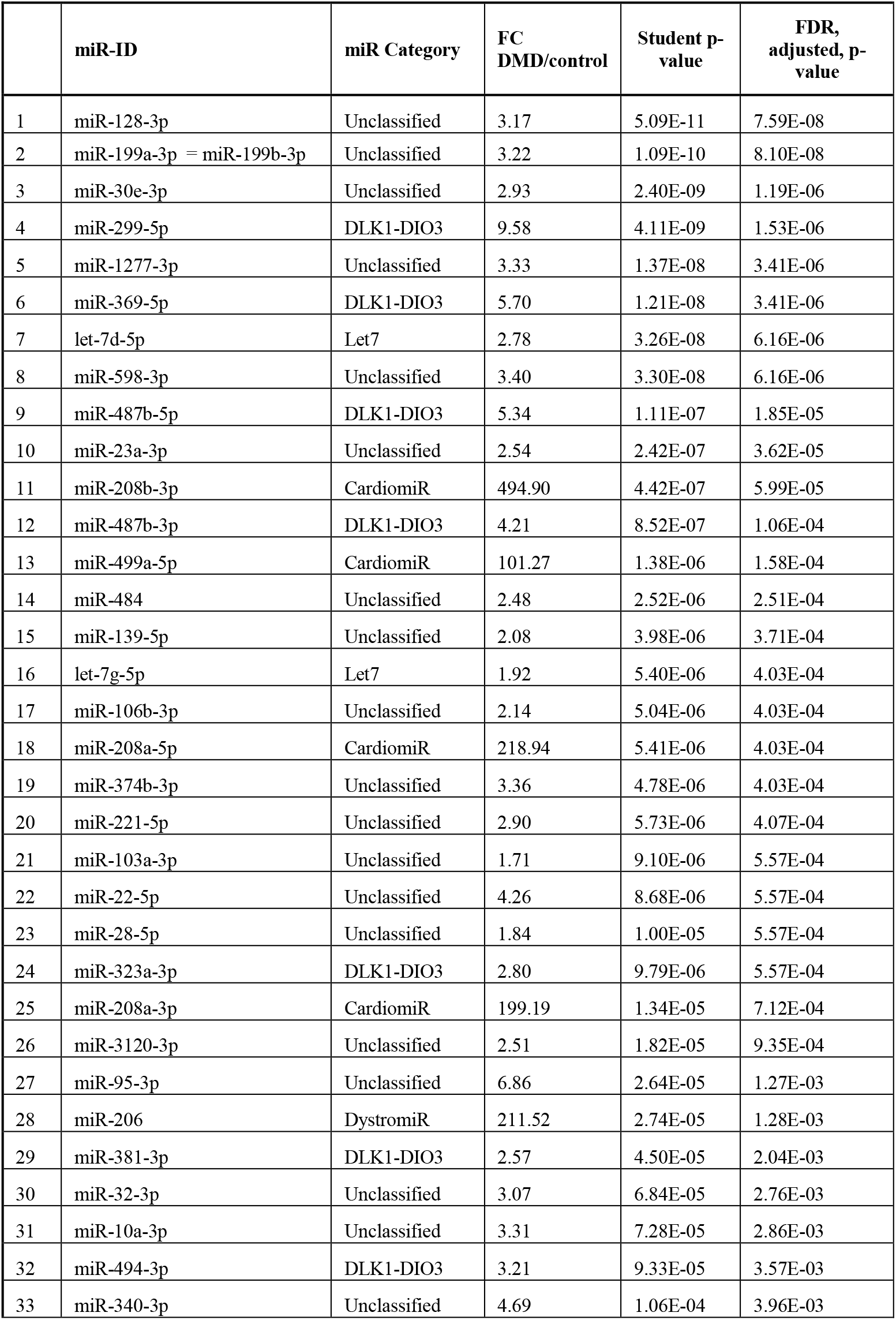
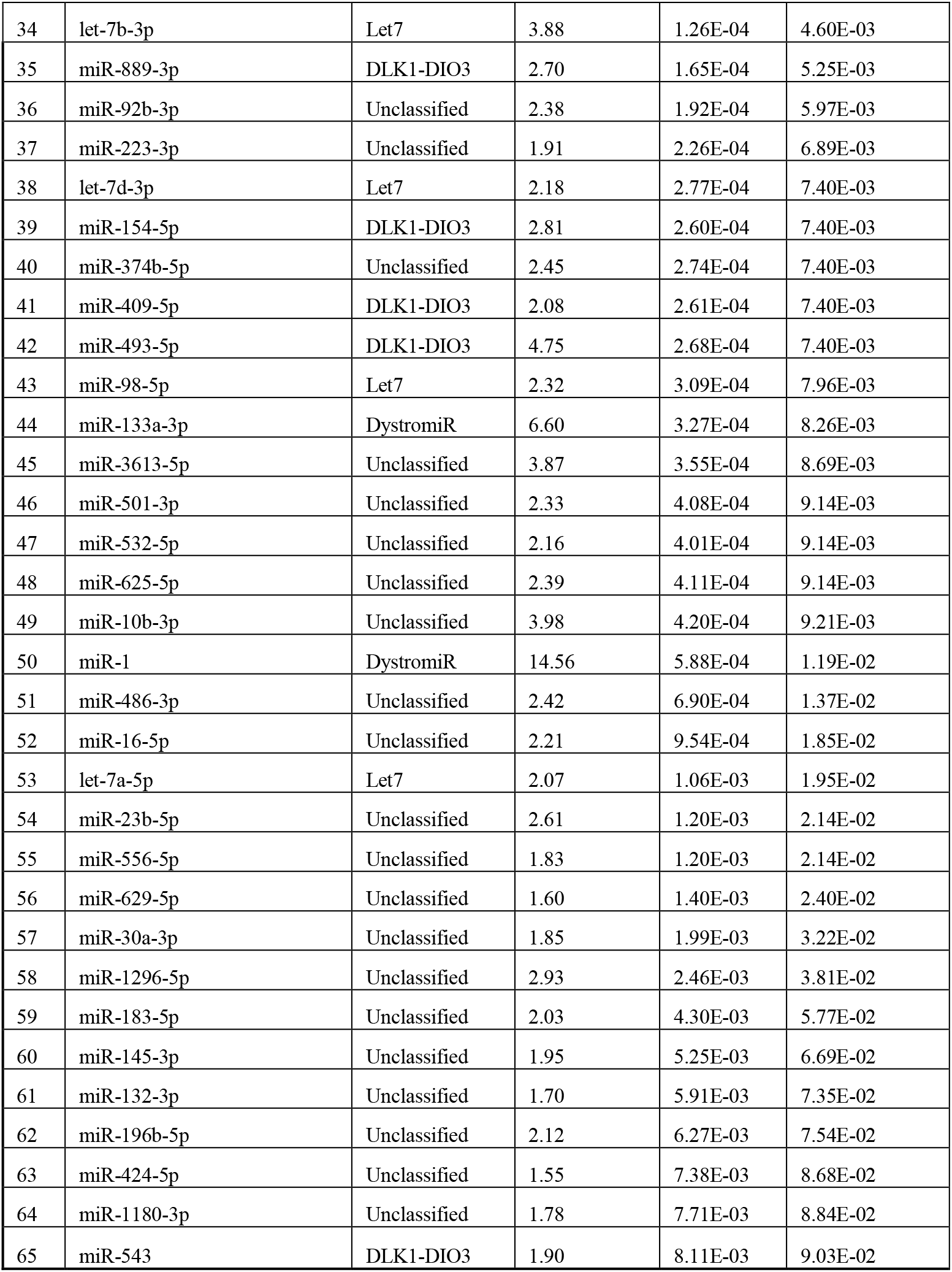
Up-regulated plasma miRNA 4-12 years old DMD versus control. MiRNA were ranked by FDR adjusted p-value. MiR categories include dystromiRs, heart-enriched miRNA (CardiomiR), miRNAs residing in the DLK1-Dio3 genomic locus, and Let-7 or miR-320 family members. MiRNAs of none of these categories were designed as unclassified.

|  | miR-ID | miR Category | FC<br>DMD/control | Student p-value | FDR,<br>adjusted, p-value |
| --- | --- | --- | --- | --- | --- |
| 1 | miR-128-3p | Unclassified | 3.17 | 5.09E-11 | 7.59E-08 |
| 2 | miR-199a-3p = miR-199b-3p | Unclassified | 3.22 | 1.09E-10 | 8.10E-08 |
| 3 | miR-30e-3p | Unclassified | 2.93 | 2.40E-09 | 1.19E-06 |
| 4 | miR-299-5p | DLK1-DIO3 | 9.58 | 4.11E-09 | 1.53E-06 |
| 5 | miR-1277-3p | Unclassified | 3.33 | 1.37E-08 | 3.41E-06 |
| 6 | miR-369-5p | DLK1-DIO3 | 5.70 | 1.21E-08 | 3.41E-06 |
| 7 | let-7d-5p | Let7 | 2.78 | 3.26E-08 | 6.16E-06 |
| 8 | miR-598-3p | Unclassified | 3.40 | 3.30E-08 | 6.16E-06 |
| 9 | miR-487b-5p | DLK1-DIO3 | 5.34 | 1.11E-07 | 1.85E-05 |
| 10 | miR-23a-3p | Unclassified | 2.54 | 2.42E-07 | 3.62E-05 |
| 11 | miR-208b-3p | CardiomiR | 494.90 | 4.42E-07 | 5.99E-05 |
| 12 | miR-487b-3p | DLK1-DIO3 | 4.21 | 8.52E-07 | 1.06E-04 |
| 13 | miR-499a-5p | CardiomiR | 101.27 | 1.38E-06 | 1.58E-04 |
| 14 | miR-484 | Unclassified | 2.48 | 2.52E-06 | 2.51E-04 |
| 15 | miR-139-5p | Unclassified | 2.08 | 3.98E-06 | 3.71E-04 |
| 16 | let-7g-5p | Let7 | 1.92 | 5.40E-06 | 4.03E-04 |
| 17 | miR-106b-3p | Unclassified | 2.14 | 5.04E-06 | 4.03E-04 |
| 18 | miR-208a-5p | CardiomiR | 218.94 | 5.41E-06 | 4.03E-04 |
| 19 | miR-374b-3p | Unclassified | 3.36 | 4.78E-06 | 4.03E-04 |
| 20 | miR-221-5p | Unclassified | 2.90 | 5.73E-06 | 4.07E-04 |
| 21 | miR-103a-3p | Unclassified | 1.71 | 9.10E-06 | 5.57E-04 |
| 22 | miR-22-5p | Unclassified | 4.26 | 8.68E-06 | 5.57E-04 |
| 23 | miR-28-5p | Unclassified | 1.84 | 1.00E-05 | 5.57E-04 |
| 24 | miR-323a-3p | DLK1-DIO3 | 2.80 | 9.79E-06 | 5.57E-04 |
| 25 | miR-208a-3p | CardiomiR | 199.19 | 1.34E-05 | 7.12E-04 |
| 26 | miR-3120-3p | Unclassified | 2.51 | 1.82E-05 | 9.35E-04 |
| 27 | miR-95-3p | Unclassified | 6.86 | 2.64E-05 | 1.27E-03 |
| 28 | miR-206 | DystromiR | 211.52 | 2.74E-05 | 1.28E-03 |
| 29 | miR-381-3p | DLK1-DIO3 | 2.57 | 4.50E-05 | 2.04E-03 |
| 30 | miR-32-3p | Unclassified | 3.07 | 6.84E-05 | 2.76E-03 |
| 31 | miR-10a-3p | Unclassified | 3.31 | 7.28E-05 | 2.86E-03 |
| 32 | miR-494-3p | DLK1-DIO3 | 3.21 | 9.33E-05 | 3.57E-03 |
| 33 | miR-340-3p | Unclassified | 4.69 | 1.06E-04 | 3.96E-03 |
| 34 | let-7b-3p | Let7 | 3.88 | 1.26E-04 | 4.60E-03 |
| 35 | miR-889-3p | DLK1-DIO3 | 2.70 | 1.65E-04 | 5.25E-03 |
| 36 | miR-92b-3p | Unclassified | 2.38 | 1.92E-04 | 5.97E-03 |
| 37 | miR-223-3p | Unclassified | 1.91 | 2.26E-04 | 6.89E-03 |
| 38 | let-7d-3p | Let7 | 2.18 | 2.77E-04 | 7.40E-03 |
| 39 | miR-154-5p | DLK1-DIO3 | 2.81 | 2.60E-04 | 7.40E-03 |
| 40 | miR-374b-5p | Unclassified | 2.45 | 2.74E-04 | 7.40E-03 |
| 41 | miR-409-5p | DLK1-DIO3 | 2.08 | 2.61E-04 | 7.40E-03 |
| 42 | miR-493-5p | DLK1-DIO3 | 4.75 | 2.68E-04 | 7.40E-03 |
| 43 | miR-98-5p | Let7 | 2.32 | 3.09E-04 | 7.96E-03 |
| 44 | miR-133a-3p | DystromiR | 6.60 | 3.27E-04 | 8.26E-03 |
| 45 | miR-3613-5p | Unclassified | 3.87 | 3.55E-04 | 8.69E-03 |
| 46 | miR-501-3p | Unclassified | 2.33 | 4.08E-04 | 9.14E-03 |
| 47 | miR-532-5p | Unclassified | 2.16 | 4.01E-04 | 9.14E-03 |
| 48 | miR-625-5p | Unclassified | 2.39 | 4.11E-04 | 9.14E-03 |
| 49 | miR-10b-3p | Unclassified | 3.98 | 4.20E-04 | 9.21E-03 |
| 50 | miR-1 | DystromiR | 14.56 | 5.88E-04 | 1.19E-02 |
| 51 | miR-486-3p | Unclassified | 2.42 | 6.90E-04 | 1.37E-02 |
| 52 | miR-16-5p | Unclassified | 2.21 | 9.54E-04 | 1.85E-02 |
| 53 | let-7a-5p | Let7 | 2.07 | 1.06E-03 | 1.95E-02 |
| 54 | miR-23b-5p | Unclassified | 2.61 | 1.20E-03 | 2.14E-02 |
| 55 | miR-556-5p | Unclassified | 1.83 | 1.20E-03 | 2.14E-02 |
| 56 | miR-629-5p | Unclassified | 1.60 | 1.40E-03 | 2.40E-02 |
| 57 | miR-30a-3p | Unclassified | 1.85 | 1.99E-03 | 3.22E-02 |
| 58 | miR-1296-5p | Unclassified | 2.93 | 2.46E-03 | 3.81E-02 |
| 59 | miR-183-5p | Unclassified | 2.03 | 4.30E-03 | 5.77E-02 |
| 60 | miR-145-3p | Unclassified | 1.95 | 5.25E-03 | 6.69E-02 |
| 61 | miR-132-3p | Unclassified | 1.70 | 5.91E-03 | 7.35E-02 |
| 62 | miR-196b-5p | Unclassified | 2.12 | 6.27E-03 | 7.54E-02 |
| 63 | miR-424-5p | Unclassified | 1.55 | 7.38E-03 | 8.68E-02 |
| 64 | miR-1180-3p | Unclassified | 1.78 | 7.71E-03 | 8.84E-02 |
| 65 | miR-543 | DLK1-DIO3 | 1.90 | 8.11E-03 | 9.03E-02 |

**Supplemental table 4.** Down-regulated plasma miRNA 4-12 years old DMD versus control. MiRNA were ranked by FDR adjusted p-value. MiR categories include the miR-320 family members and unclassified miRNAs.

|  | miR-id | miR-class | FC DMD/<br>control | Student p-value | FDR,<br>adjusted, p-value |
| --- | --- | --- | --- | --- | --- |
| 1 | miR-342-3p | Unclassified | -2.26 | 1.62E-04 | 5.25E-03 |
| 2 | miR-320a | miR-320 | -4.35 | 2.94E-04 | 7.69E-03 |
| 3 | miR-320b | miR-320 | -3.70 | 1.04E-03 | 1.95E-02 |
| 4 | miR-320d | miR-320 | -6.07 | 1.60E-03 | 2.69E-02 |
| 5 | miR-2110 | Unclassified | -2.40 | 1.95E-03 | 3.19E-02 |
| 6 | miR-361-3p | Unclassified | -3.36 | 2.23E-03 | 3.51E-02 |
| 7 | miR-320e | miR-320 | -12.69 | 2.47E-03 | 3.81E-02 |
| 8 | miR-1290 | Unclassified | -69.06 | 2.89E-03 | 4.35E-02 |
| 9 | miR-29b-3p | Unclassified | -3.64 | 3.05E-03 | 4.42E-02 |
| 10 | miR-30e-5p | Unclassified | -1.85 | 3.68E-03 | 5.18E-02 |
| 11 | miR-320c | miR-320 | -3.13 | 3.66E-03 | 5.18E-02 |
| 12 | miR-769-5p | Unclassified | -2.52 | 4.26E-03 | 5.77E-02 |
| 13 | miR-181c-3p | Unclassified | -2.78 | 5.16E-03 | 6.64E-02 |
| 14 | miR-191-5p | Unclassified | -1.70 | 5.82E-03 | 7.30E-02 |
| 15 | miR-185-5p | Unclassified | -3.67 | 6.18E-03 | 7.54E-02 |
| 16 | miR-192-5p | Unclassified | -2.92 | 6.28E-03 | 7.54E-02 |
| 17 | miR-29c-5p | Unclassified | -2.79 | 6.37E-03 | 7.54E-02 |
| 18 | miR-425-5p | Unclassified | -1.62 | 6.34E-03 | 7.54E-02 |
| 19 | miR-147b | Unclassified | -6.85 | 7.54E-03 | 8.78E-02 |
| 20 | miR-3613-3p | Unclassified | -2.32 | 7.62E-03 | 8.81E-02 |
| 21 | miR-362-5p | Unclassified | -7.44 | 7.85E-03 | 8.90E-02 |
| 22 | miR-1307-3p | Unclassified | -2.38 | 8.71E-03 | 9.56E-02 |
| 23 | miR-339-3p | Unclassified | -1.82 | 8.67E-03 | 9.56E-02 |
| 24 | miR-142-3p | Unclassified | -2.48 | 9.13E-03 | 9.75E-02 |
| 25 | miR-152-3p | Unclassified | -1.64 | 9.40E-03 | 9.88E-02 |

**Supplemental table 5:** dysregulation of glucocorticoid responsive miRNAs in 4-12 years old DMD patients. **(5a)**. Eleven DMD dysregulated miRNAs were GC-responsive. MiRNAs are classified by order of p value in treated versus untreated DMD patients. All but miR-30e-5p and miR-425-5p were upregulated in DMD and reversed by the GC treatment toward normalization. In contrast, GC-treatment increased the downregulation in DMD of miR-30e-5p and miR-425-5p. Three miRNAs, miR-27-5p, miR-379-5p and let-7i-5p were no longer dysregulated after GC treatment. Similarly, miR-27-5p, miR-379-5p were not dysregulated in the group of all DMD patients versus healthy controls (p > 0.05 in red). FC= fold change, UT = untreated, Tr = Treated. **(5b).** Heat map of miRNA dysregulation. The relative levels of the miRNAs are increased from blue to red

| miR-name | UT DMD / Healthy |  | Treated DMD /Untreated DMD |  | Treated DMD /Healthy |  | All DMD /Healthy |  |
| --- | --- | --- | --- | --- | --- | --- | --- | --- |
|  | FC | p value | FC | p value | FC | p value | FC | p value |
| miR-223-3p | 2,14 | 1,62E-05 | -1,27 | 3,15E-03 | 1,68 | 5,06E-03 | 1,91 | 2,26E-04 |
| miR-221-5p | 3,55 | 1,45E-06 | -1,57 | 3,19E-03 | 2,26 | 1,54E-03 | 2,90 | 5,73E-06 |
| miR-27b-5p | 1,52 | 3,04E-02 | -1,47 | 8,03E-03 | 1,03 | 8,89E-01 | 1,27 | 2,04E-01 |
| let-7d-5p | 3,34 | 6,32E-06 | -1,50 | 1,78E-02 | 2,22 | 1,69E-04 | 2,78 | 3,26E-08 |
| miR-28-5p | 2,08 | 3,94E-05 | -1,31 | 3,28E-02 | 1,60 | 9,26E-04 | 1,84 | 1,00E-05 |
| miR-30e-5p | -1,44 | 1,07E-02 | -1,27 | 3,55E-02 | -2,10 | 1,44E-03 | -1,85 | 3,68E-03 |
| miR-139-5p | 2,44 | 1,91E-04 | -1,41 | 4,12E-02 | 1,73 | 2,36E-04 | 2,08 | 3,98E-06 |
| let-7i-5p | 1,52 | 2,61E-03 | -1,25 | 4,36E-02 | 1,22 | 1,51E-01 | 1,37 | 1,13E-02 |
| miR-1277- | 3,89 | 3,70E-06 | -1,40 | 4,68E-02 | 2,78 | 3,97E-05 | 3,33 | 1,37E-08 |
| miR-379-5p | 1,88 | 2,19E-02 | -1,60 | 4,73E-02 | 1,18 | 5,62E-01 | 1,53 | 7,91E-02 |
| miR-425-5p | -1,44 | 2,33E-02 | -1,28 | 4,93E-02 | -1,84 | 2,40E-03 | -1,62 | 6,34E-03 |

**Supplemental table 6.** Pathway enrichment analysis of target genes for dysregulated miRNAs, considering up and down regulated miRNAs in the 4-12 years old DMD/control samples, (fdr < 0.1). Pathways are classified according p values (FDR). Shown are the 20 most enriched pathways.

|  | KEGG pathway | p (fdr) | No Genes | No miRNA |
| --- | --- | --- | --- | --- |
| 1 | Proteoglycans in cancer (hsa05205) | 1.95E-11 | 114 | 58 |
| 2 | Signaling pathways regulating pluripotency of stem cells (hsa04550) | 1.14E-10 | 88 | 54 |
| 3 | ECM-receptor interaction (hsa04512) | 1.87E-10 | 43 | 42 |
| 4 | Morphine addiction (hsa05032) | 4.29E-09 | 51 | 42 |
| 5 | Fatty acid biosynthesis (hsa00061) | 8.06E-09 | 5 | 12 |
| 6 | GABAergic synapse (hsa04727) | 6.16E-08 | 46 | 40 |
| 7 | Hippo signaling pathway (hsa04390) | 2.00E-07 | 85 | 50 |
| 8 | Hedgehog signaling pathway (hsa04340) | 1.91E-05 | 37 | 37 |
| 9 | Axon guidance (hsa04360) | 1.91E-05 | 71 | 49 |
| 10 | Cocaine addiction (hsa05030) | 2.08E-05 | 28 | 42 |
| 11 | Gap junction (hsa04540) | 2.08E-05 | 50 | 47 |
| 12 | Prostate cancer (hsa05215) | 2.08E-05 | 55 | 53 |
| 13 | Focal adhesion (hsa04510) | 2.08E-05 | 113 | 57 |
| 14 | PI3K-Akt signaling pathway (hsa04151) | 4.80E-05 | 170 | 64 |
| 15 | Estrogen signaling pathway (hsa04915) | 1.19E-04 | 53 | 55 |
| 16 | Circadian entrainment (hsa04713) | 1.39E-04 | 48 | 47 |
| 17 | FoxO signaling pathway (hsa04068) | 1.39E-04 | 75 | 52 |
| 18 | ErbB signaling pathway (hsa04012) | 1.54E-04 | 54 | 52 |
| 19 | Pathways in cancer (hsa05200) | 2.17E-04 | 191 | 58 |
| 20 | Thyroid hormone signaling pathway (hsa04919) | 2.89E-04 | 61 | 55 |

**Supplemental table 7.** List of the host genes of the identified dysregulated intragenic miRNAs that are known to be dysregulated in DMD models and patients. Both 3’ (3p) and 5’ (5p) mature miRNA forms of a given pre-miRNA were considered, when both are dysregulated. The reference column refers to the study that described the dysregulation of the host gene.

|  | miRNA name | Chrom. location | Gene symbol | Host gene Function | Knowledge of dysregulation of the host gene in dmd models and patients | Reference |
| --- | --- | --- | --- | --- | --- | --- |
| 1 | miR-128-1-3p | 2q21 | R3HDM1 | unknown | Up in dystrophin-knockdown mouse myoblast | Ghahramani Seno 2010 |
| 1* | miR-128-2-3p | 3p22 | ARPP21/R3HDM3 | unknown | Down in dystrophin-knockdown mouse myoblast |  |
| 2 | miR-1307-3p | 10q24 | USMG5/DAPIT | Mitochondrial complex 5 associated subunit | Higher in muscle biopsies of moderate versus severe DMD patients. Downregulated in dystrophic mdx muscle | Pegoraro 2011<br>Sansón 2020 |
| 3 | miR-1307-5p |  |  |  |  |  |
| 4 | miR-149-5p | 2q37 | GPC1 | Heparan sulfate proteoglycan | Up in DMD quadriceps biopsies | Fadic R 2006 |
| 5 | miR-335-3p | 7q32 | MEST | paternally imprinted mesoderm-specific transcript | miR-335 and MEST higher in mdx muscle | Hiramuki 2015 |
| 6 ** | miR-188-5p | Xp11 | CLCN5 | Chloride ion channel | Up in mdx muscle | Mizbani 2016 |
| 7 ** | miR-362-5p |  |  |  |  |  |
| 8 ** | miR-501-3p |  |  |  |  |  |
| 9 ** | miR-532-3p |  |  |  |  |  |
| 10 ** | miR-532-5p |  |  |  |  |  |
| 11 | miR-556-3p | 1q23 | NOS1AP | Nitric oxide synthase 1 adaptor protein, a cytosolic peptide that binds nNOS | Up in mdx heart and skeletal muscle | Ségalat 2005<br>Treuer 2014 |
| 12 | miR-556-5p |  |  |  |  |  |

\* The mature forms of miR-128-1-3p and miR-128-2-3p are identical and it is unknown which (or both) is dysregulated in the 4-12 DMD group. Of note, both host genes R3HDM1 and ARPP21/R3HDM3 are dysregulated in the referred dystrophic model.
\*\* These dysregulated miRNAs are clustered together at Xp11 on an intron of the CLCN5 gene

---

**Supplemental table 8.** Dysregulated intragenic miRNA in DMD plasma and their related host-gene, considering all miRNAs embedded on the sense strand of introns and exons of protein coding genes. FC and p (student T test) values are of miR expression in the blood plasma of 4-12 old DMD patients versus healthy controls. When both 3p and 5p isoforms of the same pre-miRNA were dysregulated, their host-genes was considered only once, with the lower p value miR isoform. Host genes for a number of different miRNAs were considered only once, taking into account FC and P values of the miRNA with the lowest p value.

|  | miR name | FC DMD/ control | P value | Gene symbol of host gene |
| --- | --- | --- | --- | --- |
| 1 | let-7g-5p | 1.92 | 5.40E-06 | *WDR82* |
| 2 | miR-103a-3p | 1.71 | 9.10E-06 | *PANK3* |
| 3 | miR-106b-3p | 2.14 | 5.04E-06 | *MCM7* |
| 4 | miR-107 | 1.82 | 2.55E-02 | *PANK1* |
| 5 | miR-10a-3p | 3.31 | 7.28E-05 | *HOXB3* |
| 6 | miR-10b-3p | 3.98 | 4.20E-04 | *HOXD3* |
| 7 | miR-1180-3p | 1.78 | 7.71E-03 | *B9D1* |
| 8 | miR-1228-5p | -2.73 | 9.95E-03 | *LRP1* |
| 9 | miR-126-5p | 1.67 | 2.39E-02 | *EGFL7* |
| 10 | miR-1277-3p | 3.33 | 1.37E-08 | *WDR44* |
| 11 | miR-128-1-3p | 3.17 | 5.09E-11 | *R3HDM1* |
| 12 | miR-128-2-3p | 3.17 | 5.09E-11 | *ARPP21* |
| 13 | miR-1290 | -69.06 | 2.89E-03 | *ALDH4A1* |
| 14 | miR-1294 | -3.03 | 1.40E-02 | *GALNT10* |
| 15 | miR-1296-5p | 2.93 | 2.46E-03 | *JMJD1C* |
| 16 | miR-1301-3p | -1.77 | 2.18E-02 | *DNMT3A* |
| 17 | miR-1307-3p | -2.38 | 8.71E-03 | *USMG5* |
| 18 | miR-139-5p | 2.08 | 3.98E-06 | *PDE2A* |
| 19 | miR-140-3p | -3.03 | 5.10E-02 | *WWP2* |
| 20 | miR-147b | -6.85 | 7.54E-03 | *C15orf48* |
| 21 | miR-148b-3p | -3 | 3.71E-02 | *COPZ1* |
| 22 | miR-149-5p | 2.03 | 1.49E-02 | *GPC1* |
| 23 | miR-152-3p | -1.64 | 9.40E-03 | *COPZ2* |
| 24 | miR-1537-3p | -2.86 | 1.61E-02 | *LYST* |
| 25 | miR-15b-3p | -2.08 | 2.21E-02 | *SMC4* |
| 26 | miR-185-5p | -3.67 | 6.18E-03 | *TANGO2* |
| 27 | miR-191-5p | -1.7 | 5.82E-03 | *DALRD3* |
| 28 | miR-196b-5p | 2.12 | 6.27E-03 | *HOXA9* |
| 29 | miR-2110 | -2.4 | 1.95E-03 | *C10orf118* |
| 30 | miR-23b-5p | 2.61 | 1.20E-03 | *C9orf3* |
| 31 | miR-26b-5p | 1.4 | 3.52E-02 | *CTDSP1* |
| 32 | miR-28-5p | 1.84 | 1.00E-05 | *LPP* |
| 33 | miR-30e-3p | 2.93 | 2.40E-09 | *NFYC* |
| 34 | miR-3120-3p | 2.51 | 1.82E-05 | *DNM3* |
| 35 | miR-3200-5p | -2.43 | 3.33E-02 | *OSBP2* |
| 36 | miR-320b | -3.7 | 1.04E-03 | *NVL* |
| 37 | miR-320e | -12.69 | 2.47E-03 | *PRKD2* |
| 38 | miR-32-3p | 3.07 | 6.84E-05 | *TMEM245* |
| 39 | miR-335-3p | 2.04 | 2.51E-02 | *MEST* |
| 40 | miR-339-3p | -1.82 | 8.67E-03 | *C7orf50* |
| 41 | miR-33a-5p | -5.28 | 2.06E-02 | *SREBF2* |
| 42 | miR-33b-5p | -13.52 | 5.19E-02 | *SREBF1* |
| 43 | miR-340-3p | 4.69 | 1.06E-04 | *RNF130* |
| 44 | miR-342-3p | -2.26 | 1.62E-04 | *EVL* |
| 45 | miR-361-3p | -3.36 | 2.23E-03 | *CHM* |
| 46 | miR-3615 | 1.6 | 1.53E-02 | *SLC9A3R1* |
| 47 | miR-3656 | -9.52 | 2.03E-02 | *TRAPPC4* |
| 48 | miR-378a-5p | 2.16 | 2.81E-02 | *PPARGC1B* |
| 49 | miR-4532 | -4.02 | 2.73E-02 | *NOP56* |
| 50 | miR-484 | 2.48 | 2.52E-06 | *NDE1* |
| 51 | miR-486-3p | 2.42 | 6.90E-04 | *ANK1* |
| 52 | miR-5010-5p | -2.44 | 3.97E-02 | *ATP6V0A1* |
| 53 | miR-505-5p | -1.78 | 2.87E-02 | *ATP11C* |
| 54 | miR-532-5p | 2.16 | 4.01E-04 | *CLCN5* |
| 55 | miR-548ax | 1.78 | 1.31E-02 | *ARHGAP6* |
| 56 | miR-550a-5p | -2.4 | 4.61E-02 | *ZNRF2* |
| 57 | miR-556-3p | 3.6 | 2.18E-03 | *NOS1AP* |
| 58 | miR-590-3p | 2.02 | 2.09E-02 | *EIF4H* |
| 59 | miR-598-3p | 3.4 | 3.30E-08 | *XKR6* |
| 60 | miR-625-5p | 2.39 | 4.11E-04 | *FUT8* |
| 61 | miR-627-5p | -2.69 | 1.57E-02 | *VPS39* |
| 62 | miR-628-5p | 1.97 | 2.15E-02 | *CCPG1* |
| 63 | miR-629-5p | 1.6 | 1.40E-03 | *TLE3* |
| 64 | miR-6515-5p | -2.3 | 4.15E-02 | *CALR* |
| 65 | miR-652-3p | 1.4 | 1.15E-02 | *TMEM164* |
| 66 | miR-671-3p | 1.61 | 4.23E-02 | *CHPF2* |
| 67 | miR-6852-5p | -3.54 | 1.54E-02 | *TLN1* |
| 68 | miR-7-1-3p | 1.62 | 4.53E-02 | *HNRNPK* |
| 69 | miR-7704 | -5.46 | 2.40E-02 | *HOXD1* |
| 70 | miR-95-3p | 6.86 | 2.64E-05 | *ABLIM2* |
| 71 | miR-98-5p | 2.32 | 3.09E-04 | *HUWE1* |

**Supplemental figure 1.**
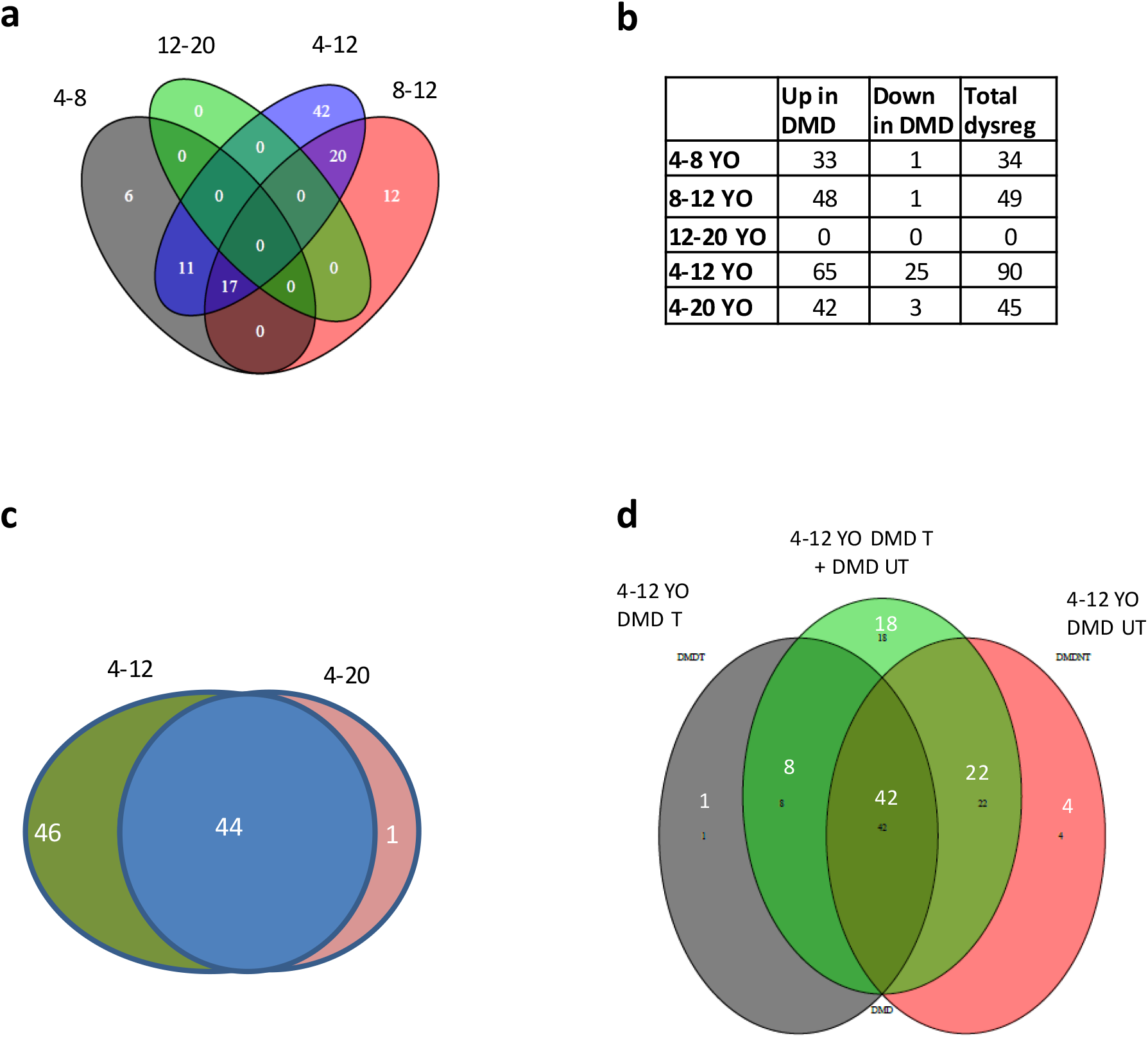
Venn diagram analysis of differentially expressed miRNAs in age and treatment stratified groups. **(a)** Venn diagram of age-specific and age overlapping dysregulated miRNAs between DMD and healthy control samples and a table presentation of (a) is shown in **(b)**. **(c)** Dysregulated miRNAs in the 4-20 years old. **(d)** Specific and overlapping dysregulated miRNAs in the 4-12 YO group between treated and untreated DMD versus healthy control. Thirty four and 49 miRNAs are dysregulated in the 4-8 and 8-12 years old group, respectively (**a and b)**. No miRNA (fdr<0.1) is differentially-expressed between DMD and healthy control in the 12-20 YO group. When age groups are combined, 90 and 45 miRNAs are dysregulated in the 4-12 and 4-20 years old groups, respectively **(a and b)**. Dysregulated miRNAs in the 4-12 years old DMD patients included all but one (miR-122) of the dysregulated miRNAs in the 4-20 years old group **(c**). When looking only at the 4-12 years old group, the classification into GC-treated and untreated as compared to the healthy control samples reduced the number of dysregulated miRNAs from 90 to 77 (**d**).

**Supplemental figure 2.**
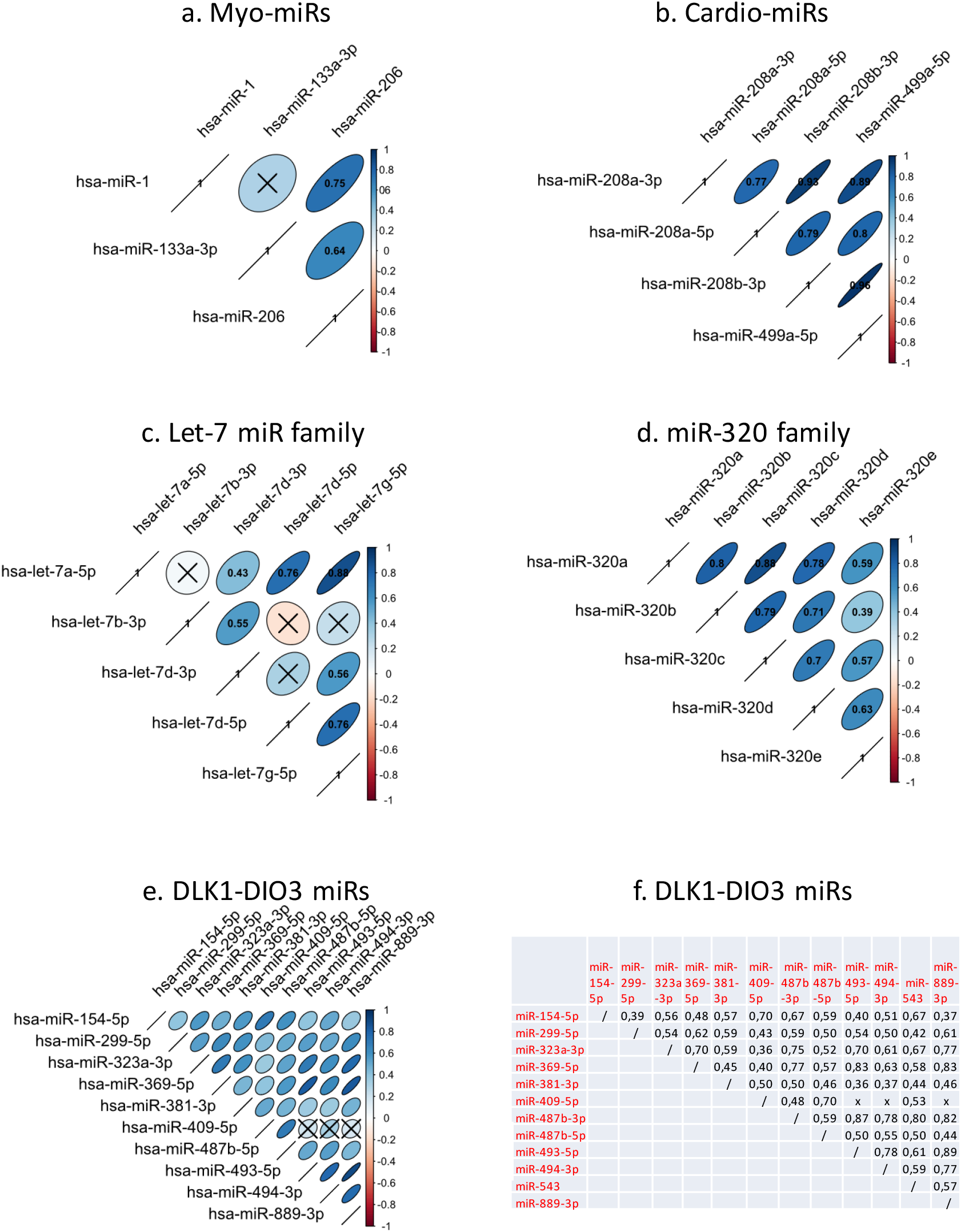
Correlogram of plasma miRNA expression in DMD patients of the 4-12 years old age group. Expression correlations were analyzed among members of miRNA classes that were identified as dysregulated in the plasma in DMD patients. Dysregulated miRNA classes included the (a) MyomiRs, (b) CardiomiR, (c) Let-7 miRs, (d) miR-320 and (e) DLK1-DIO3 cluster miRNAs. Presented only significant Spearman correlations (r <0.05) whereas “X” represent absence of correlation. The elliptic shape and the relative intensity of the blue color are proportional to the level of positive correlation. Correlation r values are presented inside the elliptic shapes. The r values for the correlations in the DLK1-DIO3 miRNAs (e) are presented in (f).

**Supplemental figure 3.**
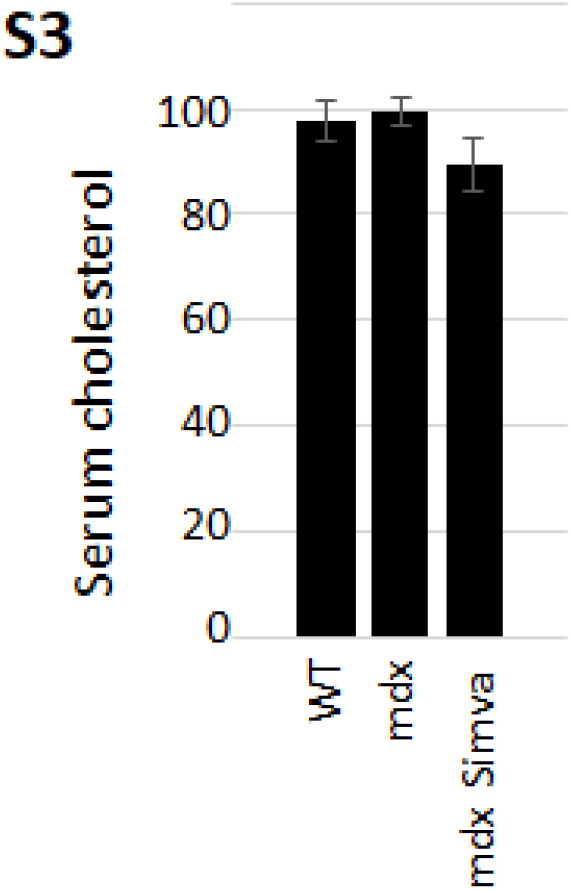
mCK and cholesterol in the serum of mdx mice treated by Simvastatin. Serum cholesterol in a 7-week old (young adult) mdx and control mice treated by Simvastatin during three weeks

**Supplemental figure 4.**
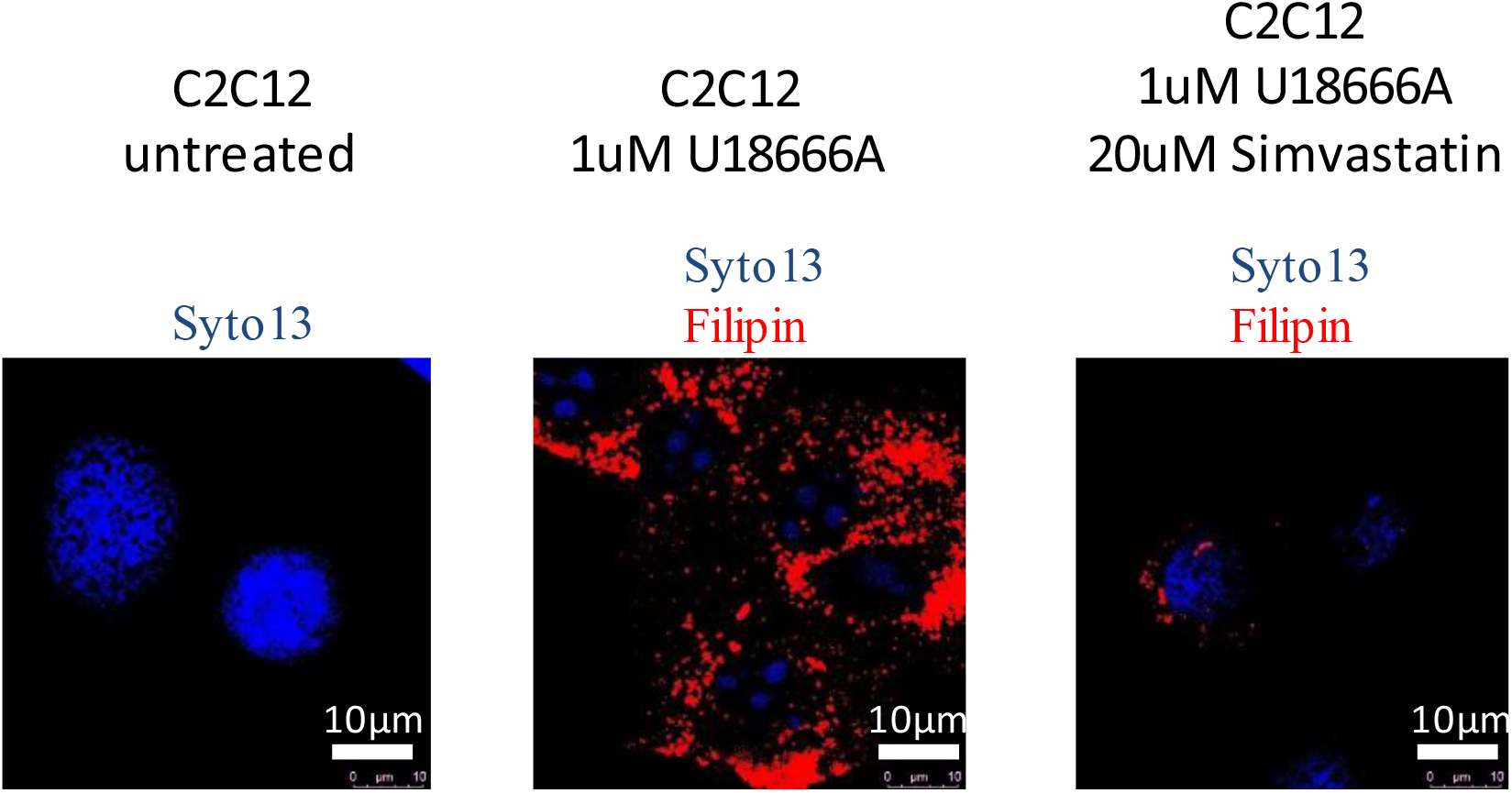
Filipin staining of free cholesterol in C2C12 cells. C2C2 myoblast were treated by the cholesterol trafficking inhibitor U18666A (Lu et al., 2015), without (middle) or with (right) 24 hours treatment by the cholesterol synthesis inhibitor, simvastatin. Nucleus were stained by Syto13. Accumulated free cholesterol was stained with filipin.

**Supplemental figure 5.**
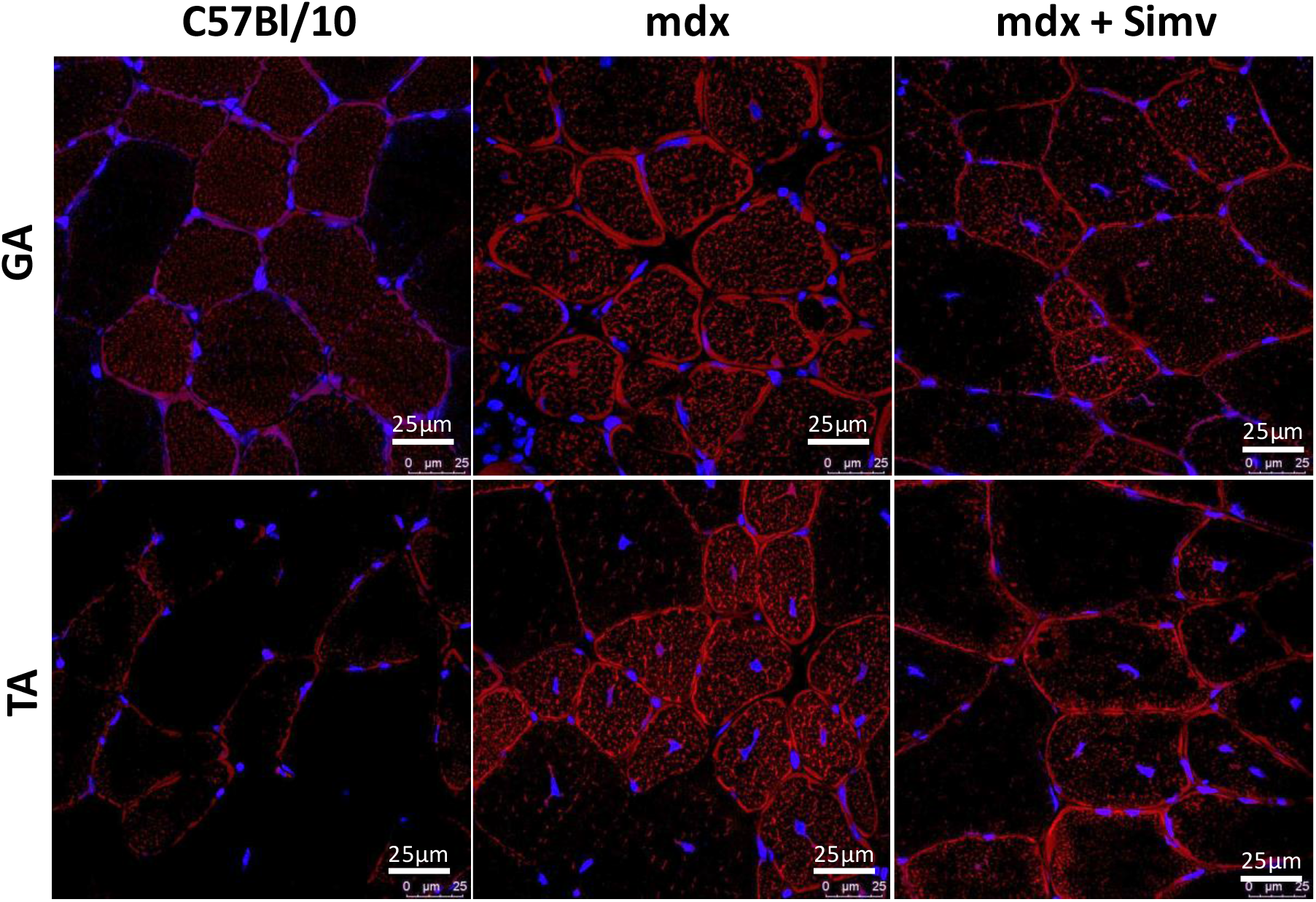
Confocal analysis of cholesterol expression in transversal sections of the Tibialis anterior and the Gastrocnemius muscles. Transversal sections of mdx and control healthy C57Bl/10 mice, Gastrocnemius (GA) and Tibialis Anterior (TA), were labeled by Cyto 13 (nucleus) in blue, and Filipins (cholesterol) in red. Mdx mice were untreated (center; mdx), or treated (right; mdx + Simva) by Simvastatin.

**Supplemental figure 6.**
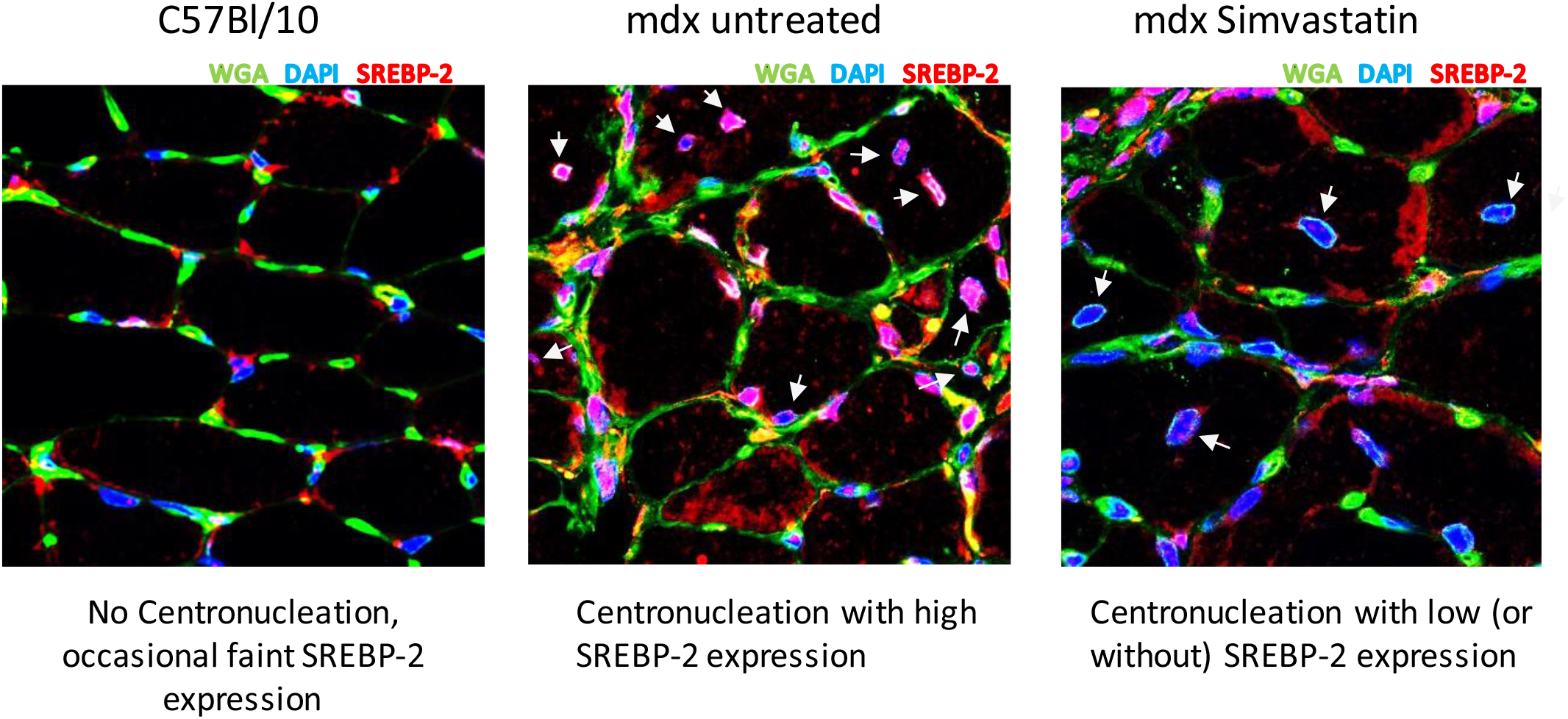
Confocal analysis of SRBP-2 expression in the mouse diaphragm. Transversal sections were immunostained with wheat germ agglutinin (WGA) in green, which stains the sarcolemma and the myonucleus membrane, with DAPI for nuclear staining, and with the anti-SREBP-2 antibody. White arrows indicating myonucleus in a central position in regenerated myofibres. Note that mdx nucleus are positive to all three colors (Pink), while only residual faint red dotes can be observed in the nucleus (blue) of the mdx treated by simvastatin, demonstrating the downregulation of nuclear SRBP-2.

**Supplemental figure 7.**
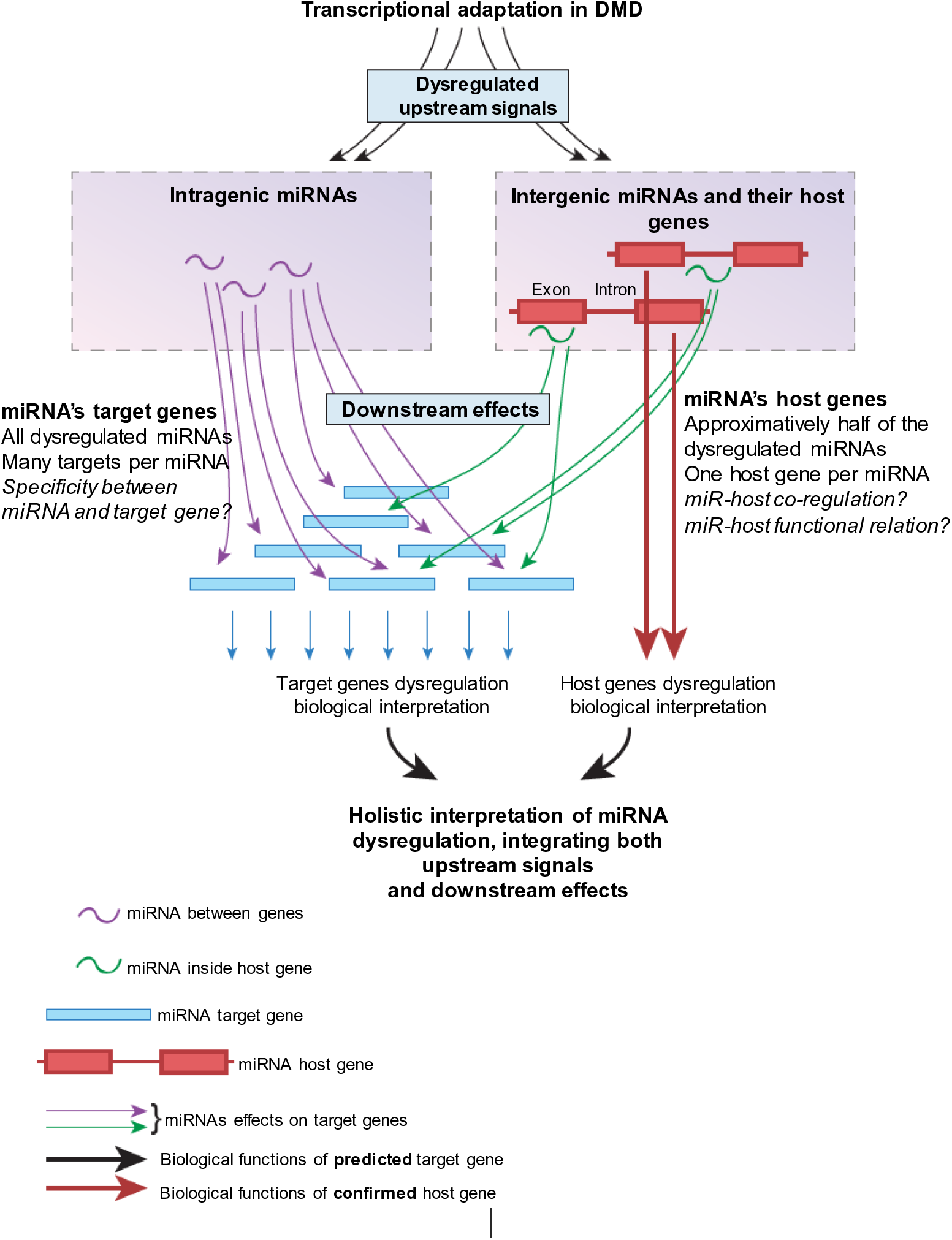
Approach for the biological interpretation of miRNA dysregulation, based on both target and host genes of the dysregulated miRNAs. Transcriptional adaptation in the diseased tissue is affecting both intergenic miRNAs (miRNA between genes, on the left), and intragenic miRNAs (miRNA inside genes, on the right). Intergenic miRNA are often co-transcribed with their host genes, cooperatively affecting downstream events. The functional link between miRNA to a host gene is thought to be strong, while the link between miRNA to their many **predicted** target gens is questionable. Dysregulation of intergenic miRNAs and their host genes provides information on **upstream signaling** in the disease, which causes transcriptional dysregulation, while the analysis of target genes may provide information on **downstream events**, which are the consequences of miRNA dysregulation.

